# Mechanisms of regulation of cryptic prophage-encoded gene products in *Escherichia coli*

**DOI:** 10.1101/2023.04.14.536235

**Authors:** Preethi T. Ragunathan, Evelyne Ng Kwan Lim, Xiangqian Ma, Eric Massé, Carin K. Vanderpool

## Abstract

The *dicBF* operon of Qin cryptic prophage in *Escherichia coli* K12 encodes the small RNA (sRNA) DicF and small protein DicB, which regulate host cell division and are toxic when overexpressed. While new functions of DicB and DicF have been identified in recent years, the mechanisms controlling the expression of the *dicBF* operon have remained unclear. Under standard laboratory growth conditions, transcription from *dicBp,* the major promoter of the *dicBF* operon, is repressed by DicA. Here, we discovered that transcription of the *dicBF* operon and processing of the polycistronic mRNA is regulated by multiple mechanisms. DicF sRNA accumulates during stationary phase and is processed from the polycistronic *dicBF* mRNA by the action of both RNase III and RNase E. DicA-mediated transcriptional repression of *dicBp* can be relieved by an antirepressor protein, Rem, encoded on the Qin prophage. Ectopic production of Rem results in cell filamentation due to strong induction of the *dicBF* operon and filamentation is mediated by DicF and DicB. Spontaneous derepression of *dicBp* occurs in a subpopulation of cells independent of the antirepressor. This phenomenon is reminiscent of the bistable switch of λ phage with DicA and DicC performing functions similar to CI and Cro, respectively. Additional experiments demonstrate stress-dependent induction of the *dicBF* operon. Collectively, our results illustrate that toxic genes encoded on cryptic prophages are subject to layered mechanisms of control, some that are derived from the ancestral phage and some that are likely later adaptations.

**Importance:** Cryptic or defective prophages have lost genes necessary to excise from the bacterial chromosome and produce phage progeny. In recent years, studies have found that cryptic prophage gene products influence diverse aspects of bacterial host cell physiology. However, to obtain a complete understanding of the relationship between cryptic prophages and the host bacterium, identification of the environmental, host or prophage-encoded factors that induce the expression of cryptic prophage genes is crucial. In this study, we examine the regulation of a cryptic prophage operon in *Escherichia coli* encoding a small RNA and a small protein that are involved in inhibiting bacterial cell division, altering host metabolism, and protecting the host bacterium from phage infections.

## Introduction

The establishment of lysogeny by temperate phages is a common occurrence in the environment, with nearly half of all sequenced bacterial genomes carrying prophages (20, 44). Whether a temperate phage will choose lysis or lysogeny depends on the metabolic state of the host cell and is driven by the activity of the phage repressor. Under specific conditions, the cognate repressor blocks transcription of the lytic genes and initiates the process of lysogeny, wherein the phage DNA is integrated into the host chromosome (20). In the absence of inducing signals, the repressor keeps a majority of phage genes transcriptionally silent in the bacterial lysogen (9). The integrated phage DNA, now called a prophage, stably replicates along with the host chromosome until a specific stress condition triggers the phage repressor to lose its control over the lytic genes, which initiates a cascade of events that drives the prophage to excise from the host chromosome and progress into the lytic cycle (14, 23).

The complex mechanisms involved in the regulation of prophage genes by the repressor and the physiological outcome of this regulation is illustrated by CI repression of λ prophage genes during lysogeny (38). CI and Cro are two repressors encoded in the immunity locus of λ. They are divergently transcribed and separated by an intergenic region that contains three operator sequences overlapping the respective promoters. For establishing and maintaining lysogeny, CI binds to two operator sequences closest to the *cro* promoter and blocks transcription of *cro*. Continuous repression by CI is necessary for stable maintenance of the prophage in the host cell. However, when the lysogen is exposed to inducing signals such as UV light or Mitomycin C, CI becomes inactivated and *cro* gene expression begins. The newly synthesized Cro binds to the operators overlapping the *cI* promoter sequence and in turn, blocks transcription of *cI*. With *cI* transcription blocked, the prophage genes necessary to initiate a lytic cycle are expressed and new λ progeny phages are produced (15, 37, 38). This genetic control of lytic and lysogenic cycles by two repressors is almost universally conserved in lambdoid phages, with some phages having additional layers of regulation (15).

Qin is one of the four lambdoid cryptic prophages of *Escherichia coli* K12 and lacks the majority of the replication, head, and tail genes (13). Of particular interest in this prophage is the *dicBF* operon which encodes the small RNA DicF and small protein DicB. DicB interacts with MinC and targets it to the nascent septum at cell center, resulting in MinC-dependent depolymerization of FtsZ, resulting in cell division inhibition and filamentation of *E*. *coli* (24, 25). We recently showed that the small protein DicB confers an advantage to the host cell by providing superinfection immunity against certain phages. This protection is specific to phages that use the ManYZ inner membrane proteins of the Mannose PTS to inject their DNA into the host cell (40). A second regulator in the *dicBF* operon, the small RNA DicF, base pairs with and inhibits *ftsZ* mRNA translation, limiting FtsZ protein synthesis (4). We found previously that DicF also directly affects host cell metabolism by inhibiting translation of pyruvate kinase, xylose regulator, and mannose transporter mRNAs (3, 4). These studies on DicB and DicF demonstrate how prophage-encoded regulators perform diverse functions, some of them beneficial, in the host cell.

The promoter of the *dicBF* operon, *dicBp*, is similar to the λ P_L_ promoter and is repressed by the DicA repressor (5, 6). The *dicAC* locus, located immediately upstream of the *dicBF* operon, is similar in arrangement and sequence to the immunity locus of lambdoid phages. DicA is analogous to the P22 C2 repressor and DicC to the P22 Cro repressor (6). Interestingly, DicA differs from the conventional lambdoid repressors in lacking the alanyl-glycyl bond necessary for RecA-mediated cleavage during the SOS response (6). The conditions leading to derepression of *dicBp* and production of DicB and DicF are still unknown.

In this study, we characterized the transcriptional and post-transcriptional mechanisms of regulation of the *dicBF* operon. During stationary phase, we observed that the sRNA DicF accumulates and is processed from the *dicBF* mRNA transcript by RNase III and RNase E. Characterization of transcriptional regulation of *dicBp,* the major promoter of the *dicBF* operon, revealed that DicA repression of *dicBp* is relieved by Rem, a putative antirepressor protein encoded on the Qin cryptic prophage. We show that the Rem antirepressor promotes filamentation of cells due to induced expression of the cell division inhibitors DicB and DicF. Our results also demonstrate that spontaneous induction of *dicBp* occurs in a subpopulation of *E*. *coli* cells when the *dicBF* operon is deleted and that stress conditions including urea and high temperature can also induce transcription from *dicBp*. Overall, this study identifies multiple distinct mechanisms by which the activity of an unconventional repressor of a lambdoid cryptic prophage is regulated and the consequences of derepression leading to the production of prophage-encoded products that influence various physiological processes of the host bacterium.

## Results

### DicA represses transcription of the *dicBF* operon

The *dicBF* operon of Qin cryptic prophage, which encodes the small RNA DicF and small protein DicB, is highly conserved in many strains of *E*. *coli* (4). The *dicBF* operon, which also encodes *ydfA*, *B*, *C*, *D*, and *E*, is under the control of the regulators encoded by the *dicAC* locus, located immediately upstream of the operon (Fig. 1A) (6). Previous studies have identified DicA as the repressor of the *dicBF* operon (6, 7, 47). The *dicAC* locus is similar to the immunity locus of lambdoid phages with DicA similar to the CI repressor and DicC similar to Cro repressor (6). The intergenic regions between *dicA* and *dicC*, and *dicA* and *dicBp*, contain three operator sequences similar to those controlling λ P_R_ and P_L_ promoters, respectively (6, 47).

**Figure 1.**
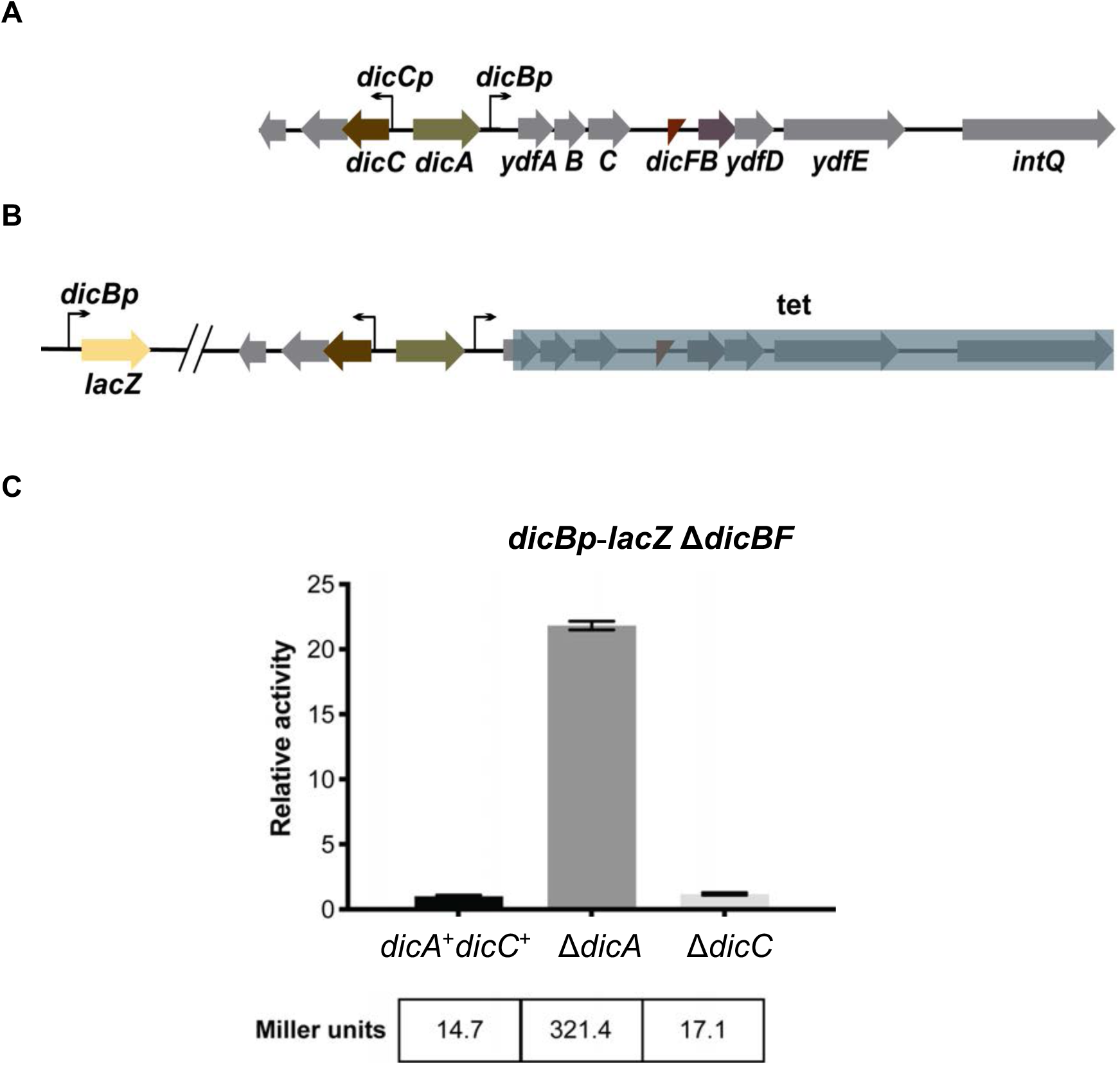
Regulation of the *dicBF* operon by DicA. A) The *dicAC* locus is located immediately upstream of the *dicBF* operon and resembles the immunity locus of lambdoid phages. The promoter of the *dicBF* operon, *dicBp*, is repressed by DicA. B) Reporter strain PR221 contains an out-of-locus *dicBp*-*lacZ* fusion, and the *dicBF* operon genes and *intQ* are deleted and replaced with a tetracycline resistance marker (*dicBp*-*lacZ* Δ*dicBF*). C) β-galactosidase activity of the *dicBp*-*lacZ* Δ*dicBF* strain was assayed with deletion of *dicA* and *dicC*. The specific activities in Miller units (indicated at the bottom) were normalized to the control strain (*dicA*^+^*dicC*^+^) to obtain the relative activity for each experimental strain. Error bars were calculated as standard deviation from three biological replicates.

To confirm the roles of DicA and DicC in regulation of *dicBp* in our strain background, we constructed a transcriptional fusion with *dicBp* fused to *lacZ* placed at a locus distal to the Qin prophage (Fig. 1B). Next, we deleted the *dicBF* operon and *intQ* in this strain and replaced it with a tetracycline resistance marker (hereafter called *dicBp*-*lacZ* Δ*dicBF*), which allows us to study the regulation of *dicBp* without the growth inhibitory effect caused by induction of the *dicBF* genes (Fig. 1B) (4). We tested the roles of DicA and DicC in regulating *dicBp* by deleting the corresponding genes in the *dicBp*-*lacZ* transcriptional fusion strain and carrying out β-galactosidase assays (Fig. 1C). We found that deletion of *dicA* induced *dicBp* expression by 22-fold compared to the control strain. Deletion of *dicC* did not affect *dicBp* activity (Fig. 1C). These results support the previous observations (6, 47) that DicA is the repressor of the *dicBF* operon.

### The sRNA DicF of the *dicBF* operon is expressed during stationary phase growth

The *dicBF* polycistronic transcript initiates upstream of *ydfA* and encompasses 6 protein coding sequences (*ydfABC, dicB, ydfDE*) and *dicF*, which codes for the DicF sRNA (Fig. 1A). Prior work suggested that DicF production requires processing of the longer polycistronic mRNA that initiates upstream of *ydfA* (19), but one study suggested that there is another promoter upstream of *dicB* (12). To understand the conditions and regulatory mechanisms governing expression of the *dicBF* operon, we first examined how DicF is produced.

The sRNA DicF of the *dicBF* operon is one of the few cryptic prophage-encoded sRNAs whose relevance in the host bacterium has been established (4, 33, 42). To identify conditions that lead to the production of DicF in *E*. *coli* K12, we tracked levels of this sRNA during growth in different media. RNA was extracted at various timepoints from wild-type (WT) *E*. *coli* K12 MG1655 growing aerobically either in LB medium or M63 medium supplemented with glucose and iron (Fe). DicF levels were monitored by Northern Blot. Work by Faubladier, *et al*., had shown that alternative processing of the DicF 5’ end by RNase III and RNase E generates a long 190 nucleotides (nt) species and a short 53 nt species, respectively (19). In our growth conditions, we were able to detect DicF fragments of both 190 nt and 53 nt lengths in LB and M63 minimal medium supplemented with glucose and FeSO_4_ (Fig. 2A, B, C, D). Levels of both DicF species were low during log phase growth but increased and remained stable during stationary phase in the two media used (Fig. 2A, B). Together, these results indicate that DicF is produced during stationary phase when *E*. *coli* is growing aerobically.

**Figure 2.**
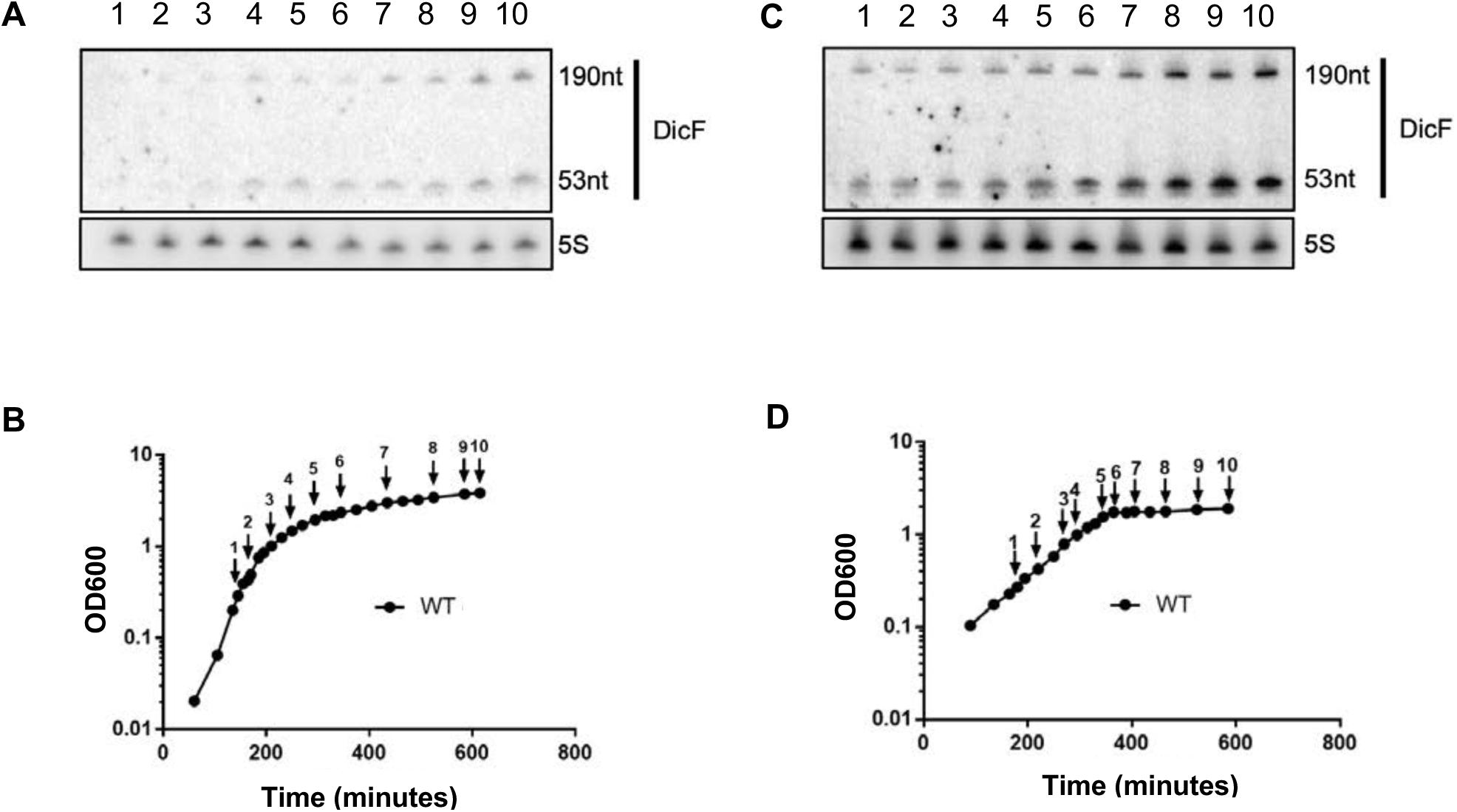
Production of the sRNA DicF. A) Northern blots showing DicF levels in *E*. *coli* K12 MG1655 growing in LB medium. 5S RNA was used as a loading control. B) Growth curve *E. coli* MG1655 in LB medium showing time points for RNA extraction. Numbers correspond to lanes in A. C) Northern blots showing DicF levels in *E. coli* MG1655 grown in M63 minimal medium supplemented with 0.2% glucose and 1µM FeSO_4_. 5S RNA was used as a loading control. D) Growth curve *E. coli* MG1655 in M63 minimal medium supplemented with 0.2% glucose and 1µM FeSO_4_ showing time points for RNA extraction. Numbers correspond to lanes in C.

Next, we investigated the mechanism by which the sRNA DicF was processed from the *dicBF* mRNA transcript during the stationary phase of *E*. *coli* growth. RNase III and RNase E have been implicated in previous studies to yield the 5’ end of the functional DicF RNA from the longer transcript originating from *dicBp* (Fig. 3A and (19)). Using a strain with thermosensitive RNase E (TS) (30), we observed that the 190-nt DicF RNA fragment increased in abundance at 43°C, when RNase E (TS) is inactivated, while the 53-nt fragment was not detected (Fig. 3B). However, the 190-nt DicF fragment disappeared when RNase III was absent. Together, these data establish that a functional RNase E is required to generate the minimal 53nt DicF fragment and RNase III cleavage is necessary for generating the longer 190nt DicF fragment, supporting the previously suggested processing pattern of DicF (19). Primer extension analysis demonstrated that the 5’ end of the 190-nt DicF is generated by RNase III cleavage between positions 813 and 814 (Fig. S1A, B) consistent with previous observations (19) and suggesting that there is no additional promoter within the *dicBF* operon. Thus, both RNase III and RNase E play vital roles in transcript processing to generate the sRNA DicF.

**Figure 3.**
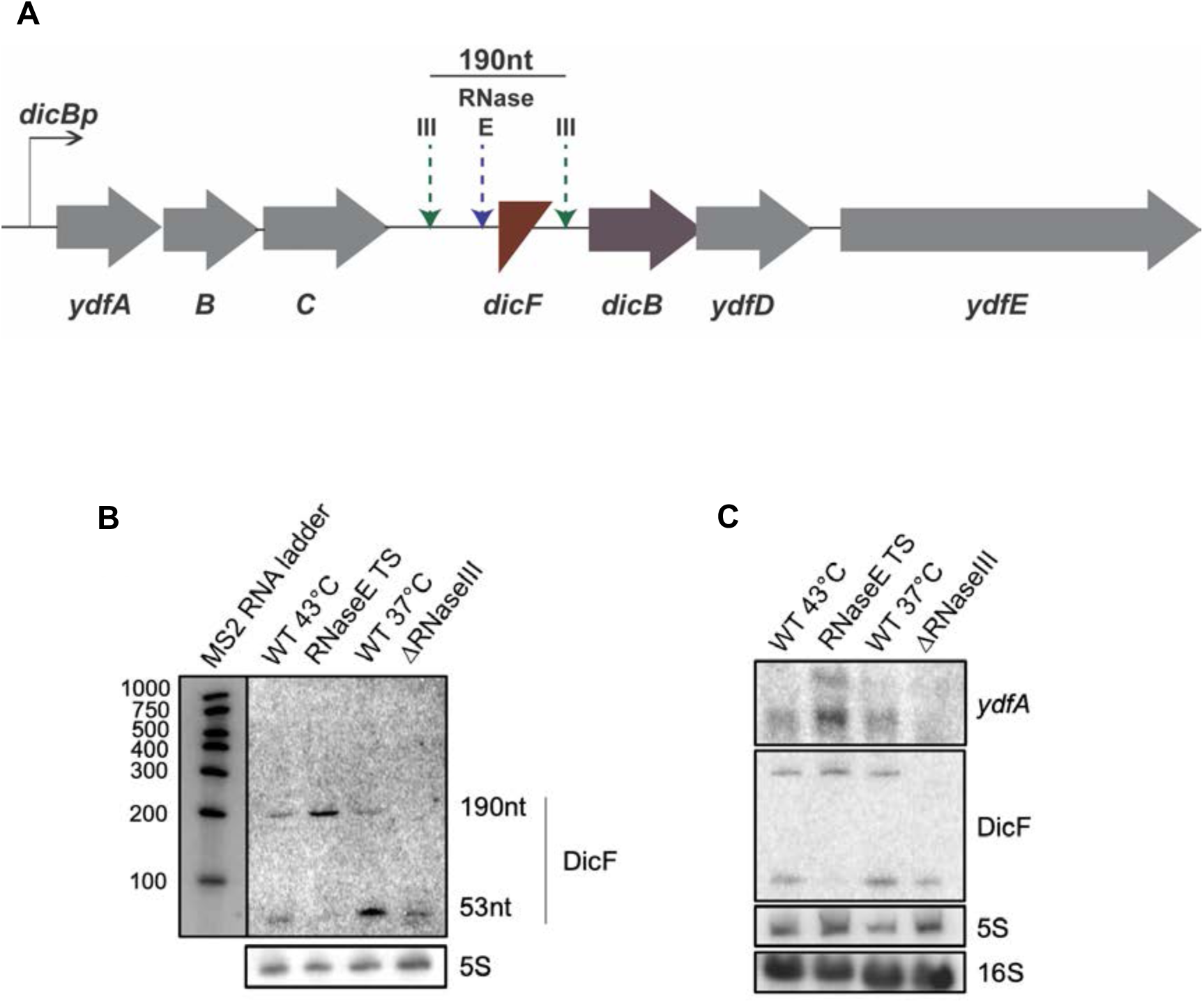
DicF is generated by processing of a transcript originating at *dicBp* by RNase III and RNase E. A) RNase III and RNase E have been implicated in the processing of DicF from the *dicBF* transcript at the sites indicated (19). B) Northern blot showing DicF levels in strains grown in M63 minimal medium supplemented with 0.2% glucose and 1µM FeSO_4_. RNase E TS is thermosensitive and was inactivated by heat shock at 43°C for 15 minutes. MS2 ladder was used to determine each fragment’s length. 5S RNA is used as a loading control. C) A Northern blot using probes against *ydfA* mRNA and DicF was performed using RNA extracted from overnight cultures of indicated strains grown in M63 minimal medium supplemented with 0.2% glucose and 1µM FeSO_4_. RNase E TS was inactivated as described in B. 5S and 16S RNA were used as loading controls.

We established that DicF was generated from the polycistronic transcript initiating at *dicBp* by probing for *ydfA* mRNA (Fig. 3C). We observed *ydfA* transcripts in cells grown to stationary phase, the same condition where we observed the highest levels of DicF (Fig. 3C, 2D). The *ydfA* transcript was observed in WT cells at both 43°C and 37°C and was more abundant in the RNase E (TS) strain at the non-permissive temperature (Fig. 3C), suggesting that RNase E-dependent processing upstream of *dicF* destabilizes the upstream region of the polycistronic transcript. We infer based on the patterns of accumulation and locations of processing that the lower *ydfA* band corresponds to the portion of the transcript encompassing *ydfABC*, while the upper band that accumulates only in the RNase E (TS) strain corresponds to *ydfABC-dicF*. We did not detect any longer species that would correspond to *ydfABC-dicFB-ydfDE.* In the Δ*rnc* strain lacking RNase III, we did not detect either *ydfA* band, suggesting that RNase III-dependent processing may somehow stabilize the upstream portion of the polycistronic transcript. Finally, by performing RT-PCR on RNA extracted from WT cells grown in minimal medium, we demonstrated that transcripts encompassing the entire region between *ydfA-dicF* are detected in WT cells (Fig. S2A, B). Together, these data strongly suggest that DicF is generated by RNase E- and RNase III-mediated processing from a polycistronic transcript originating from *dicBp*. Since we find no evidence for other promoters in this region, we proceeded to further study transcriptional regulation at *dicBp*.

### An antirepressor protein derepresses the *dicBF* operon

DicA strongly represses *dicBp* under laboratory growth conditions ((6), Fig. 1C). DicA is predicted to bind to operator sequences that overlap the promoter and exclude RNA polymerase binding, similar to other lambdoid repressors (6). However, DicA differs from the conventional P22 or λ repressor because it is significantly shorter in length (135 amino acids compared to 216 amino acids of P22 C2 repressor) and lacks the alanyl-glycyl bond that is necessary for RecA-mediated cleavage during SOS response (6). This suggests that its activity may be regulated by a different mechanism than the conventional repressors. In a study by Lemire, *et al*., (26), DicA was identified as one of the prophage repressors in *E*. *coli* that was similar in sequence to the Gifsy prophage repressors GfoR and GftR in *Salmonella enterica* serovar Typhimurium. These repressors also share other common features such as having shorter length than the conventional lambdoid repressors and lacking the alanyl-glycyl bond. Instead, Gifsy repressor activity was found to be regulated by antirepressor proteins, encoded either on the same prophage or another prophage harbored in the same strain (26). Due to the similarity of DicA to the Gifsy repressors, we sought to identify an antirepressor protein of DicA in *E*. *coli* K12.

Using the protein sequence of GfoA, the antirepressor of Gifsy-1 prophage repressor, we performed a Position-Specific Iterative (PSI)-BLAST search to find similar proteins in *E*. *coli*. Since many protein hits generated in the first round of PSI-BLAST did not have identifiable homologs in *E*. *coli* K12, we performed a subsequent homology search using protein hits that had comparable length to GfoA. These secondary searches identified Rem, encoded by a gene on Qin prophage 3.6 kb upstream from the *dicBF* operon. Other potential antirepressors were identified on other cryptic prophages in the MG1655 genome. These included YpjJ of CP4-57 prophage, YeeT of Cp4-44 prophage, and YkfH of CP4-6 prophage.

To test whether any of the putative antirepressors impacted induction of the *dicBF* operon, the genes encoding putative antirepressor proteins were placed under the control of an inducible (P*_tet_*) promoter and a heterologous ribosome binding site (RBS) for expression from a plasmid. These were transformed into a strain with an in-locus transcriptional *dicBp*-*lacZ* fusion to monitor transcription. Notably, because this is an in-locus fusion, the *lacZ* insertion effectively makes the fusion strain null for the cell division inhibitors DicB and DicF. Production of the predicted antirepressors was induced and Miller assays were carried out to quantify the β-galactosidase activity. Compared to the control strain, the strain expressing *rem* had a 165-fold increase in *dicBp* expression (Fig. 4). Strains producing the other putative antirepressors had *lacZ* activity similar to the control strain (Fig. 4). We confirmed that expression of *rem* from a construct with its native RBS also strongly induces the *dicBp* promoter (Fig. S3A). A third construct where *rem* was placed under the control of a P*_lac_* promoter also resulted in derepression of *dicBp*-*lacZ* in a Δ*dicBF* strain (Fig. S3B). These results suggest that the cryptic prophage Qin encodes the Rem antirepressor that antagonizes the activity of DicA, resulting in derepression of *dicBp*.

**Figure 4.**
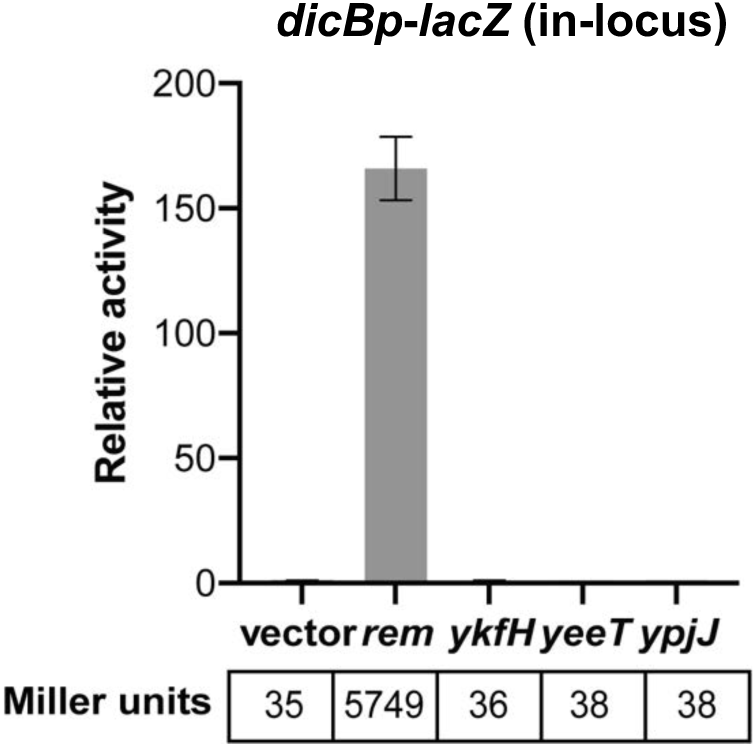
Rem acts as an antirepressor of *dicBp*. The predicted antirepressors were cloned on plasmids under P*_tet_* control and introduced to a strain carrying a *dicBp*-*lacZ* in-locus transcriptional fusion. The antirepressor genes were induced for three hours with 10 ng/ml anhydrous tetracycline and β-galactosidase activity was assayed. The relative activity was calculated by dividing the Miller units of the specific strain to that of vector control. Error bars were calculated as standard deviation from three biological replicates.

### Rem induces filamentation of *E*. *coli* cells

With the identification of Rem as the antirepressor of the *dicBF* operon, we wanted to examine Rem-dependent phenotypes and determine whether these were related to expression of the *dicBF* operon. Cells filament when the *dicBF* operon is expressed due to the combined activities of the cell division inhibitors DicB and DicF. For this experiment, *rem* was expressed in the WT strain, a *dicB* mutant, a *dicF* mutant, a *dicB dicF* double mutant, and a *qin* mutant (where the entire Qin prophage was deleted). The WT strain expressing *rem* was highly filamentous compared to the same strain harboring the vector control (Fig. 5A). Expression of *rem* in the Δ*qin* background did not result in cell filamentation, suggesting that Qin-encoded gene products were responsible for the filamentous phenotype (Fig. 5A). Strains with individual deletions of *dicB* or *dicF* were filamentous when *rem* was expressed. However, in the Δ*dicB*Δ*dicF* strain, *rem* expression did not lead to filamentation as cells appeared similar to the WT with vector control. These results indicate that when *dicBp* is induced by a Rem-dependent mechanism, production of either DicB or DicF is sufficient to inhibit cell division. In strains expressing the other three putative antirepressors, we did not observe filamentation (Fig. S4A).

**Figure 5.**
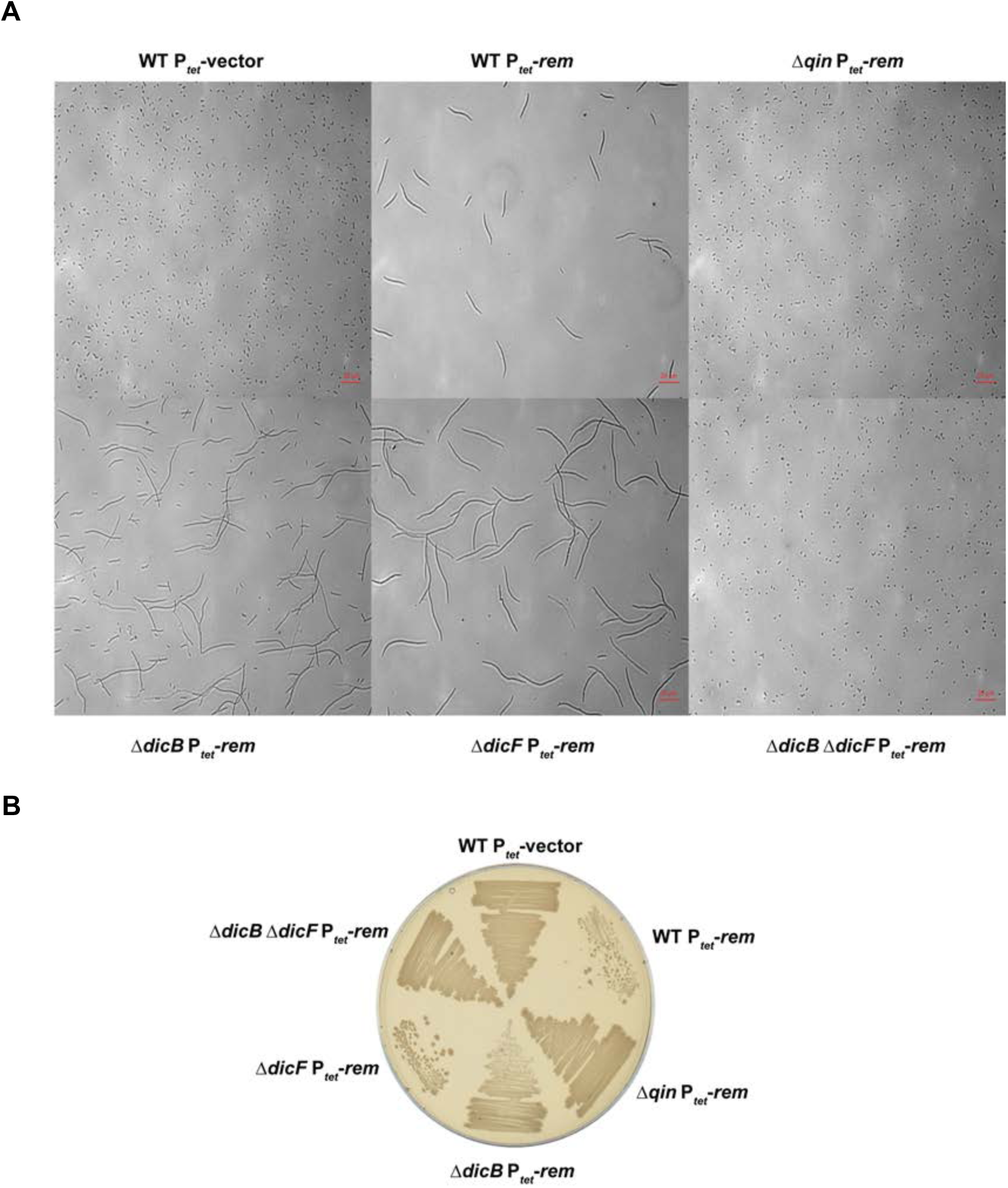
Expression of *rem* leads to filamentation and growth inhibition of cells. A) WT strain and strains with deletions of *dicB*, *dicF*, *dicB dicF*, and the entire *qin* cryptic prophage harboring P*_tet_*-vector or P*_tet_*-*rem* plasmids were grown for three hours with 100 ng/ml anhydrous tetracycline and observed under a microscope. B) The strains used in A were streaked on LB agar plates with 100 ng/ml anhydrous tetracycline and incubated overnight at 37°C.

Growth of strains expressing *rem* yielded results consistent with the microscopy. The WT strain expressing *rem* was severely growth inhibited compared to the vector control strain, whereas in the Δ*qin* strain, Rem overproduction did not inhibit growth (Fig. 5B). The Δ*dicF rem-*expressing strain was more growth inhibited than the Δ*dicB rem-*expressing strain, consistent with our previous study indicating that the small protein DicB is a more potent growth inhibitor than the sRNA DicF (4). The Δ*dicB*Δ*dicF* strain was not growth inhibited and looked similar to the vector control strain (Fig. 5B). We did not observe growth inhibition when any of the other putative antirepressors were overproduced (Fig. S4B). Collectively, our results are consistent with the model that Rem acts as an antirepressor of the *dicBF* operon and expression of *rem* specifically derepresses *dicBp* leading to production of the cell division inhibitors DicF and DicB.

We tested whether deletion of *rem* influences DicF levels during growth in minimal medium during stationary phase. Deletion of *rem* did not substantially affect accumulation of DicF (Fig. S5), suggesting that induction of the *dicBF* operon can occur by an antirepressor-independent mechanism.

### *dicBp* switches on spontaneously in a subpopulation of cells

Under laboratory conditions, *dicBp* is repressed by DicA (Fig. 1C and (6)). However, when an overnight culture of *dicBp*-*lacZ* Δ*dicBF* strain (Fig. 1B) was diluted and plated on LB agar plates with X-Gal (40 µg/ml), we observed growth of a few blue colonies among a background of white colonies. The blue colonies indicate that these colonies induced *dicBp*-*lacZ* to a high level. This induction of *dicBp* in a subset of cells is reminiscent of the spontaneous induction of λ lysogens, which is a well-characterized phenomenon that occurs at low frequency (38, 43).

To further characterize the spontaneous induction phenomenon, *dicBp*-*lacZ* Δ*dicBF* strains that carried different deletions in *qin* prophage genes or the host factor *recA* were constructed (Table 1). We made dilutions of overnight cultures in phosphate-buffered saline (PBS) and plated reporter strains on LB agar with X-Gal plates. The frequency of colonies that showed high-level *dicBp* activity was quantified by counting the number of blue (promoter on) and white (promoter off) colonies. In the *dicBp*-*lacZ* strain with deletion of the *dicBF* locus, 1.16% of colonies were blue, demonstrating that ∼1 out of every 100 cells in an overnight culture had switched on *dicBp* (Table 1).

**Table 1.** dicBp spontaneously induced in a sub-population of cells. Overnight cultures of dicBp-lacZ strains were diluted in PBS and plated on LB agar plates with X-Gal (40 µg/ml). The percentage of colonies with dicBp induced was calculated using (number of blue colonies/total number of blue and white colonies)*100. The standard error was calculated as the standard deviation of values from three biological replicates.

| Strains (*dicBp-lacZ*) | Genotype | % of *dicBp*-on colonies | Standard error |
| --- | --- | --- | --- |
| PR219 | *dicBF*+ | 0 | 0 |
| PR221 | $\Delta dicBF$ | 1.16 | 0.55 |
| PR231 | $\Delta dicBF \Delta dicA$ | 100 | 0 |
| PR232 | $\Delta dicBF \Delta dicC$ | 0.04 | 0.07 |
| XM252 | $\Delta dicBF \Delta rem$ | 1.67 | 0.75 |
| XM260 | $\Delta dicBF \Delta recA$ | 1.02 | 0.54 |

Deletion of *dicA* resulted in 100% blue colonies since the absence of DicA constitutively turns on *dicBp* (Table 1 and Fig. 1C). Deletion of the Cro-like *dicC* reduced the frequency of blue colonies to 0.04%. In phage λ, Cro repressor is responsible for flipping the bistable genetic switch by repressing *cI* transcription in a lysogen, which causes induction of lytic genes from P_L_ (38). In the absence of *cro*, P_L_ induction (due to derepression by CI) is not sustained, leading to lower levels of expression of lytic genes encoded by the λ operon that is analogous to the *dicBF* operon (6, 38). By analogy to the CI-Cro genetic switch, in strains lacking DicC, spontaneous induction of *dicBp*, would not result in DicC-mediated repression of *dicA* transcription to reinforce the induction of *dicBF* operon transcription. This would result in restoration of DicA-mediated repression of *dicBp* and yield the lower frequency of cells that stably induce *dicBp* in the Δ*dicC* strain (Table 1). Together, these results suggest that DicA and DicC of the Qin cryptic prophage constitute a functional bistable genetic switch.

Spontaneous induction of λ prophage occurs at a frequency of 1 in 10^5^ lysogens (28). SOS induction in a small population of cells was determined to play an important role in the spontaneous induction of λ lysogens, as a RecA deletion mutant and a CI mutant that does not get cleaved yielded reduced phage production from spontaneous induction (22, 35, 38). To check if this spontaneous induction of *dicBp* is also mediated by RecA, we deleted *recA* in *dicBp*-*lacZ* Δ*dicBF* strain and found that frequency of spontaneous induction was 1.02% (Table 1), similar to the *recA*^+^ strain. This implies that the observed *dicBp* derepression is independent of RecA activity. Next, we deleted the antirepressor *rem* in *dicBp*-*lacZ* Δ*dicBF* and quantified the frequency of *dicBp* induction. The frequency was 1.67%, which is similar to that of the *rem^+^* strain (Table 1). Thus, the observed mechanism of *dicBp* spontaneous induction is also independent of the antirepressor.

Interestingly, we did not observe any blue colonies in the *dicBF^+^ dicBp*-*lacZ* strain (Table 1). This is likely because high-level induction of *dicBp* would be toxic to the cells in this strain because both DicB and DicF are potent cell division inhibitors (4). Thus, *dicBF^+^* cells that induced *dicBp* in overnight culture would likely not form colonies. An alternative hypothesis is that a gene product encoded in the *dicBF* operon could contribute to maintaining tight DicA-mediated repression of *dicBp*.

### *dicBp* is induced by urea and high temperature

A study in *E. coli* K12 MG1655 showed that DicF-dependent filamentation occurred under anaerobic conditions, due to increased stability of DicF under anaerobic conditions and faster degradation under aerobic conditions (36). Another study showed that under microaerobic conditions, four DicF orthologs encoded by different prophages in *E*. *coli* O157:H7 are produced (33). However, we have not observed DicB or DicF-mediated filamentation of MG1655 cells under anaerobic conditions or increased expression of DicF under microaerobic conditions (data not shown). We also tested the expression of *dicBp*-*lacZ* in response to the SOS-inducing agent Mitomycin C by disk diffusion assay on indicator plates and observed that, unlike λ and other lambdoid prophages, mitomycin C does not induce *dicBp* (data not shown).

To identify other conditions that induce expression of the *dicBF* operon, we carried out a screen using Biolog plates (10). We grew the *dicBp*-*lacZ* Δ*dicBF* strain in fourteen different 96-well Biolog phenotype microarray plates. Plates contained different carbon sources, antibiotics and other chemicals, or compounds that induced different osmotic and ionic effects and pH changes. X-Gal was added to plates to monitor LacZ activity. These assays identified urea as an inducer of *dicBp-lacZ*. To further characterize induction of *dicBp* by urea, we streaked Δ*dicBF* strains harboring *dicBp*-*lacZ* on LB X-Gal plates with different concentrations of urea (Fig. 6). We observed that *dicBp*-*lacZ* Δ*dicBF* colonies were white on plates with no urea but turned blue when urea was present, with colonies turning darker blue in increasing concentrations of urea up to 2% (Fig. 6). The *dicBp-lacZ* Δ*dicBF* Δ*rem* and *dicBp-lacZ ΔdicBF* Δ*recA* strains were induced to similar levels as the *dicBp-lacZ ΔdicBF* parent strain, indicating that *dicBp* induction by urea was independent of the antirepressor Rem and the SOS response (requiring RecA) (Fig. 6). The *dicBp-lacZ ΔdicBF* Δ*dicA* strain was blue with and without urea as expected since it lacks the repressor. We observed that *dicBp-lacZ ΔdicBF* Δ*dicC* colonies turned only a light blue color on 2% urea plates and was not as highly induced as the *dicBp-lacZ ΔdicBF* parent (Fig. 6). This suggests that the induction of *dicBp* by urea could occur via flipping of the bistable switch between DicA-mediated repression of *dicBp* to DicC-mediated repression of *dicA*, as seen previously with the spontaneous induction phenotype. Finally, we did not observe *dicBp-lacZ dicBF+* colonies turning blue in the presence of urea.

**Figure 6.**
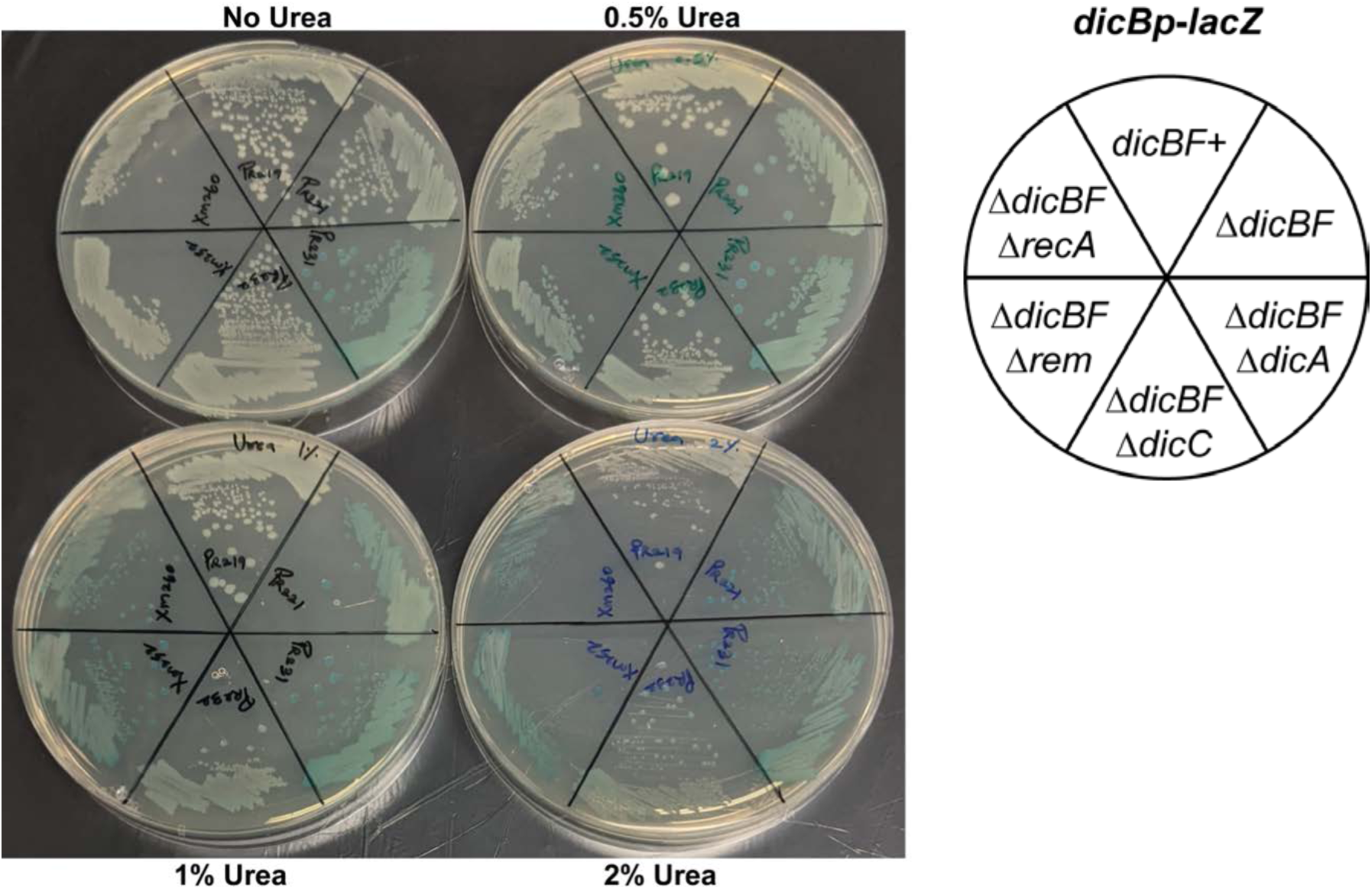
*dicBp* is induced by urea. *dicBp*-*lacZ* strains with different deletions of the *dic* locus, *rem*, and *recA* were streaked on LB agar plates with X-Gal (40 µg/ml) and urea at the indicated concentrations and incubated overnight at 37°C.

We hypothesized that induction of *dicBp* by urea might be due to DicA instability because urea is a protein denaturing agent. Since high temperature is another physiological condition that destabilizes proteins, we tested *dicBp* expression in our reporter strains at different temperatures. At 30°C and 37°C, all strains except *dicBp*-*lacZ* Δ*dicBF* Δ*dicA* were white on LB X-Gal plates (Fig. 7A). At 39°C, *dicBp*-*lacZ* Δ*dicBF*, *dicBp*-*lacZ* Δ*dicBF* Δ*rem*, and *dicBp*-*lacZ* Δ*dicBF* Δ*recA* turned light blue. At 42°C, the colonies of these strains turned a darker blue (Fig. 7A). This shows that DicA-mediated repression of *dicBp* is relieved at 39 and 42°C in strains lacking *dicBF*, and this effect was independent of Rem and RecA. As observed with urea, *dicBp*-*lacZ* Δ*dicBF* Δ*dicC* turned only a light blue at 42°C (Fig. 7A). The *dicBp*-*lacZ dicBF*+ colonies turned a very light blue only at 42°C (Fig. 7A). In β-galactosidase assays performed using liquid cultures, all strains except *dicBp*-*lacZ* Δ*dicBF* Δ*dicA* had very low β-galactosidase activity at 30°C and 37°C (Fig. 7B). The β-galactosidase activity of *dicBp*-*lacZ* Δ*dicBF* Δ*dicA* remained high at all the three temperatures tested since the repressor is deleted in this strain. At 42°C, the β-galactosidase activity of the remaining strains increased 3- to 4-fold compared to the activity of the same strain at 30°C (Fig. 7B). It was interesting to note that *dicBp*-*lacZ dicBF*+ and *dicBp*-*lacZ* Δ*dicBF* Δ*dicC* also had a similar increase in β-galactosidase activity at 42°C compared to the respective strains at 30°C, which was somewhat different than what we observed on the X-Gal plates (Fig. 7A, B). Another important observation was that the β-galactosidase activity of *dicBp*-*lacZ* Δ*dicBF* strain was approximately twice that of *dicBp*-*lacZ dicBF*+ strain at all three temperatures (Fig. 7B). Overall, this suggests that *dicBp* is derepressed at higher temperature and the presence of the wild-type *dicBF* operon contributes to the repressive effect on *dicBp*.

**Figure 7.**
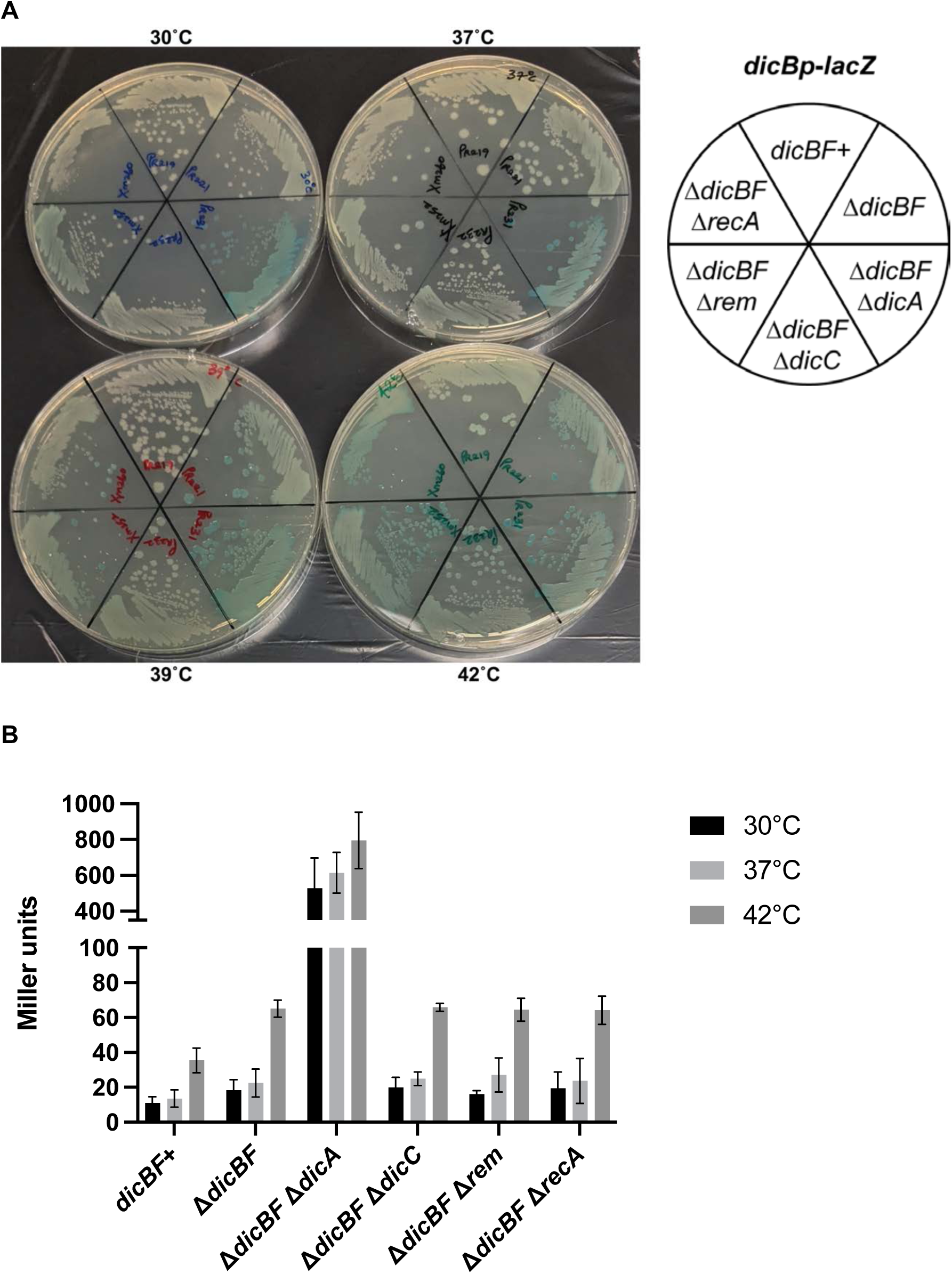
*dicBp* is induced by high temperature. A) The indicated strains were streaked on LB agar plates with X-Gal (40 µg/ml) and incubated overnight at 30, 37, 39, or 42°C. B) The indicated strains were cultured at 30, 37, or 42°C from overnight cultures grown at 30°C. When the strains reached an OD_600_ of ∼0.8, β-galactosidase activity was assayed. Error bars were calculated as standard deviation from three biological replicates.

## Discussion

The roles of the cryptic prophage products DicB and DicF in protecting host cells from phage infections (40), altering host metabolism (4), and inhibiting cell division (4, 5, 11, 24) indicate the complex relationship that exists between the host cell and the Qin cryptic prophage (Fig. 8A). The evolutionary maintenance of these regulators hints at possible fitness advantages that DicB and DicF confer on the host cell under specific conditions. Protection of host cells from phage infections by DicB is a clear example of such a beneficial relationship (40). Other work has implicated cryptic prophages (including Qin) and cryptic prophage products (including DicB) in protecting bacteria from stressors such as sublethal concentrations of antibiotics (45). In this study, we investigated the mechanisms by which production of DicB and DicF are regulated and found several levels of control that appear to modulate the stability or activity of the prophage repressor protein DicA (Fig. 8A, B). Under standard laboratory growth conditions, DicA represses the *dicBF* operon and *dicC*, resulting in low levels of expression of DicB and DicF (Fig. 8B). Nevertheless, we can detect expression of the *dicBF* operon from *dicBp* leading to production of DicF via RNase E- and RNase III-dependent processing of the *dicBF* polycistronic mRNA (Fig. 3 and (19). DicF accumulates particularly in stationary phase (Fig. 2). We identified an induced state wherein DicA-mediated repression of the *dicBF* operon is abrogated by an antirepressor protein Rem (Fig. 8B), which is encoded on the same cryptic prophage (Fig. 4). Ectopic production of Rem leads to derepression of the *dicBF* operon, resulting in production of DicB and DicF and causing cell filamentation (Fig. 4, 5). We do not yet know what signal stimulates the Rem-dependent fully induced state. Production of DicF during the stationary phase of growth is independent of Rem (Fig. S5), suggesting that there is at least one additional mechanism for induction of the *dicBF* operon. Spontaneous induction of *dicBp*, in a subpopulation (∼1%) of cells is also independent of the Rem antirepressor protein (Fig. 8B, Table 1). The spontaneous induction appears to occur by a mechanism that is reminiscent of the phage λ bistable switch. We showed that transcription of the *dicBF* operon can be induced by external factors like urea and high temperature via a Rem-independent mechanism which also influences the bistable switch of DicA and DicC. (Figs. 6, 7). Collectively, we uncovered multiple conditions and mechanisms that impact expression of the *dicBF* operon of Qin cryptic prophage.

**Figure 8.**
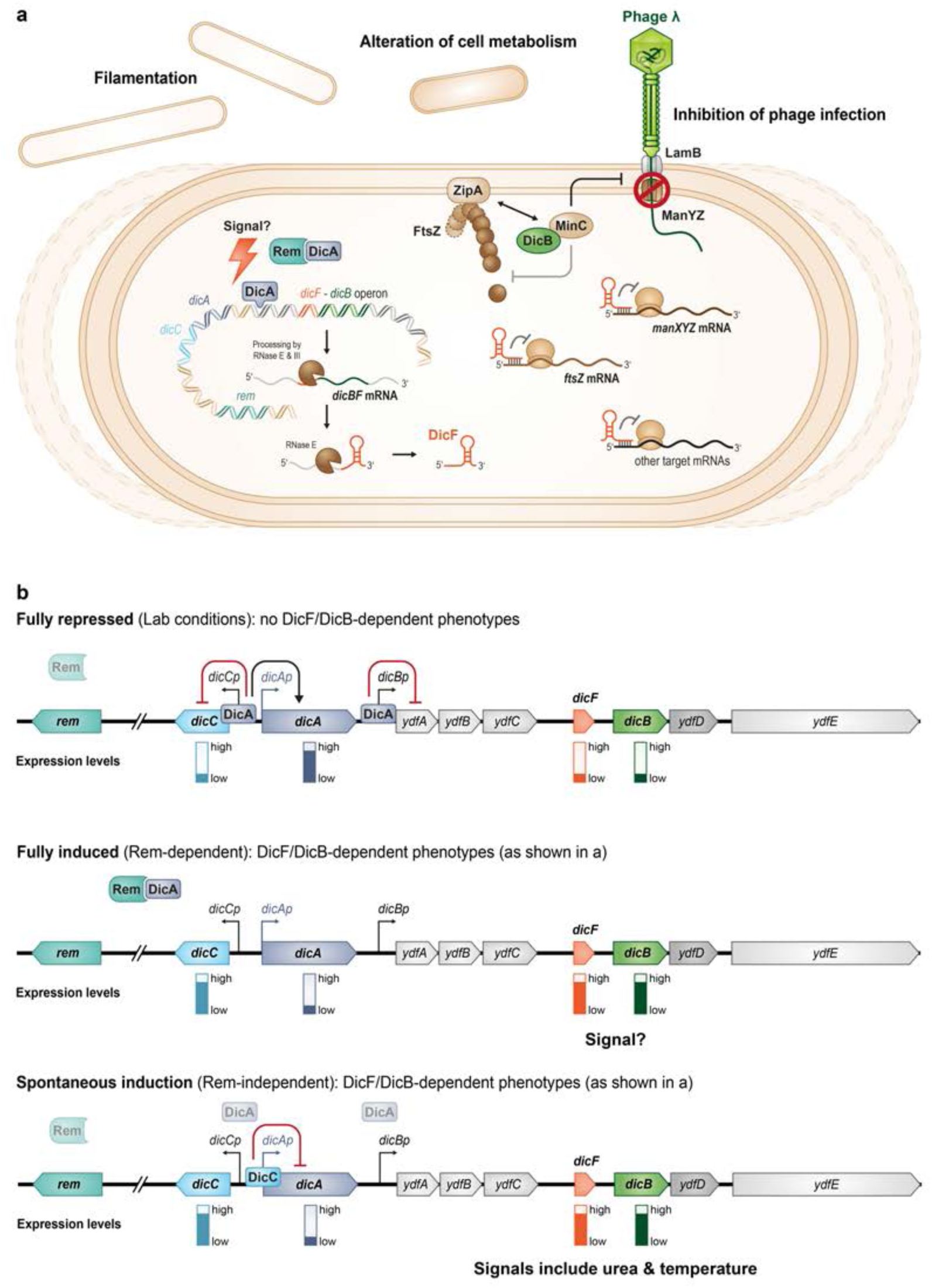
Working model for regulation of the *dicBF* operon. **(A)** The sRNA DicF and the small protein DicB are encoded on the Qin prophage of *E*. *coli* K12. In this study, we showed that DicF accumulates during stationary phase and is generated from the polycistronic mRNA by RNase E- and RNase III-dependent processing. Previous work demonstrated that DicF basepairs with and prevents translation of *ftsZ* and *manXYZ* mRNAs resulting in inhibition of cell division and alteration of carbohydrate metabolism, respectively. DicB affects FtsZ and ManXYZ protein activity. DicB localizes MinC, a negative regulator of FtsZ polymerization, to the septum to inhibit cell division. The DicB-MinC complex also inhibits ManXYZ function and prevents transport of mannose and infection by phages that require ManXYZ for DNA injection. **(B)** The *dicBF* operon is regulated by DicA and DicC, which are analogous to λ phage CI and Cro, respectively, and encoded upstream of *dicBp*. DicA represses transcription of the *dicBF* operon and *dicC*. Transcription at *dicBp* is regulated by several mechanisms. The antirepressor Rem eliminates the repressive effect of DicA on *dicBp* and induces production of DicF and DicB, resulting in filamentation. *dicBp* can also be induced spontaneously in a subset of cells. DicC is important for this phenotype, which resembles the λ bistable switch. Environmental signals like urea and high temperature can impact the bistable switch and induce transcription from *dicBp* in a DicC-dependent manner.

The induction of a functional prophage in a lysogen is primarily dependent on repressor inactivation. Representatives of a major class of repressors, like the λ prophage repressor CI, get directly cleaved during the SOS response, which results in expression of the lytic genes that were under its control (38). However, there is another class of repressors that do not get cleaved during the SOS response. The Gifsy prophages of *Salmonella* encode the repressors GfoR and GfhR that fall into this category (26). The regulation of these repressors involve antirepressor proteins that directly interact with and disassociate the cognate repressor from the operator sequence, resulting in transcription from the derepressed promoter (26). These antirepressors are under the control of LexA, which responds to SOS-inducing conditions (26). It is interesting that a cryptic prophage like Qin has an intact regulation module consisting of the repressor and the antirepressor. Given the multiple functions of DicB and DicF in the host cell, and the conservation of the *dicBF* operon in many *E*. *coli* strains (4, 40), expression of the *dicBF* operon could be beneficial for the host under specific conditions. Unlike the Gifsy antirepressors, the condition that induces antirepressor Rem production is not known. Remarkably, a functional temperate phage, named mEp460_evo81 (32), was recently isolated from the virome of infant feces and it harbors genes similar to *dicB*, *dicF*, *dicAC* and *rem*. Our analyses indicate that DicA and Rem of mEp460_evo81 share similarity with *E*. *coli* K12 DicA and Rem in size and amino acid sequence. The conservation of DicA and Rem in a functional phage suggests that the regulation of DicA by the antirepressor Rem is important for the lifecycle of the phage and that signals that are specific to the environment that the host bacterium resides in could trigger derepression of the *dicBF* operon through Rem.

A recent study suggested that one additional mechanism of regulation of the *dicBF* operon involves repression by the nucleoid associated proteins Hfq and Fis. Combining mutations in *hfq* and *fis* is lethal in *E*. *coli* K12, but growth is partially rescued by the deletion of cryptic prophages Qin or DLP-12 (2). Both Qin and DLP-12 contain intact phage lysis cassettes that can lyse the host cell when expressed. Additionally, Qin harbors DicB, DicF, and HokD, three gene products whose prolonged expression can directly affect bacterial growth and survival (4, 21). We speculate that host proteins Hfq and Fis could be involved in regulating the immunity regions of these cryptic prophages to silence prophage genes that can be toxic to the host cell when overexpressed. Since the repressor of the Qin prophage is regulated by an antirepressor protein, these nucleoid associated proteins could also directly repress the transcription of *rem* to inhibit antirepressor-mediated induction of *dicBp*.

The bistable switch of λ controls complex genetic circuits involved in the decision between the lysogenic or lytic lifecycle. CI and Cro are the two repressors involved in this genetic switch. The role of CI is to block expression of genes involved in the lytic cycle by binding to the operator sequences at P_R_ (analogous to the *dicCp*) and P_L_ (analogous to *dicBp*) (Fig. 8B) (38). CI binding to P_R_ prevents transcription of *cro*, while also activating its own transcription. Under stress conditions that lead to inactivation of CI, Cro is produced, which in turn blocks *cI* expression. This depletion of CI leads to the expression of λ lytic genes and the prophage commits to the lytic cycle (38, 41). DicA and DicC are similar to λ CI and Cro in terms of their structural similarity and binding to operator sequences in *dicCp* (P_R_) and *dicBp* (P_L_) promoters (6). A study by Yun *et al*. in *E*. *coli* MG1655 showed that by binding to *dicCp*, DicA blocks *dicC* transcription and activates its own transcription (47). Recent work in *E*. *coli* O157:H7 showed that overexpression of DicC repressed *dicA*’s transcription, similar to Cro repression of *cI* (39). Additionally, deletion of *dicC* led to a reduction in *dicB* levels while overexpression induced *dicB* expression (39). Our results show that DicC is important for the spontaneous induction phenotype of *dicBp*, as deletion of *dicC* results in a reduction in the frequency of cells that have induced *dicBp*. DicC is also necessary for a strong induction of *dicBp* with urea and high temperature. Thus, along with supporting data from previous studies, our results are consistent with a model where the cryptic prophage repressors DicA and DicC form a functional bistable switch in *E*. *coli* K12, and an inherent instability of DicA causes derepression of *dicBp* and initiates the lytic cycle equivalent of Qin in a subpopulation of cells (Fig. 8B).

The cryptic prophages of the model bacterium *E*. *coli* K12 were shown to increase resistance of the host to environmental stresses, like oxidative stress and osmotic stress, and to certain antibiotics, like quinolones and β-lactams (45). Our lab’s work on DicF and DicB of the *dicBF* operon of Qin cryptic prophage revealed the influence of these prophage-encoded products on metabolism (3, 4), cell division (4), and phage susceptibility of *E*. *coli* K12 (40). Since prophages also encode gene products like lytic proteins and toxins that can have undesired effects on the host cell viability, it is common for such prophage genes to be strongly repressed in the host bacterium. So far, very few studies have investigated the mechanisms that lead to derepression of the repressors of *E*. *coli* K12 cryptic prophages, especially those of unconventional repressors like DicA. The *E*. *coli* K12 cryptic prophage Rac also encodes an unconventional repressor, called RacR, that shares similarity with the Gifsy prophage repressors and do not respond to SOS response (26). Remarkably, Rac also encodes a Cro-like protein called YdaS (14), suggesting that the bistable switch of λ phage CI-Cro could exist in yet another cryptic prophage of *E*. *coli* K12. Based on the findings from the current study, it would be interesting to explore if cryptic prophages with DicA-like unconventional repressors in *E*. *coli* K12 and other bacteria encode the associated antirepressors to regulate the expression of genes under repressor control. Exploring the relationship between the conditions that lead to the expression of cryptic prophage genes and functions of the associated gene products in the host will provide further understanding on the complex role of cryptic prophages in host bacteria.

## Materials and Methods

### Strain construction

All the strains and phages used in this study are summarized in Table 2 and the oligonucleotides (from Integrated DNA Technologies) are listed in Table 3. The strains used in the study are derivatives of *E*. *coli* K12 strain MG1655. Chromosomal mutations were constructed using the λ red recombination method as described previously (16, 17, 46).

**Table 2.** Strains and plasmids used in this study.

| Strain name | Genotype | Source |
| --- | --- | --- |
| DB166 | DJ480 *lacIq* TetR SpecR | This study |
| DB181 | DJ480 *lacIq* TetR SpecR Δ*dsrA*::tet | This study |
| EM1055 | DJ480 (MG1655 Δ*lacX174*) | (31) |
| EM1277 | EM1055 *rne-3071 zce-726*::Tn10 | (30) |
| EM1321 | EM1055 *rnc-14*::Tn10 | (30) |
| ENG63 | EM1055 Δ*dicF*::kan | This study |
| ENG262 | EM1055 Δ*rem*::cat | This study |
| PM1805 | DJ480 P<sub>BAD</sub>::*cat-sacB*::*lacZ*, mini λTet | (29) |
| PR136 | DB166 *dicBp-lacZ* (in-locus) | This study |
| PR142 | DB166 Δ*qin*::kan | This study |
| PR143 | DB166 Δ*dicB*::kan | This study |
| PR144 | DB166 Δ*dicF*::kan | This study |
| PR145 | DB166 Δ*qin*::scar | This study |
| PR146 | DB166 Δ*dicB*::scar | This study |
| PR147 | DB166 Δ*dicF* ::scar | This study |
| PR148 | PR147 Δ*dicB*::scar | This study |
| PR149 | DB166 Δ*dicF* Δ*dicB* | This study |
| PR219 | PM1805 *dicBp-lacZ* | This study |
| PR221 | PM1805 *dicBp-lacZ* $\Delta$(*ydfA-intQ*)::tet | This study |
| PR226 | PR221 Δ*dicA*::cm | This study |
| PR227 | PR221 Δ*dicC*::cm | This study |
| PR231 | PR221 Δ*dicA*::scar | This study |
| PR232 | PR221 Δ*dicC*::scar | This study |
| XM252 | PR221 Δ*rem*::scar | This study |
| XM260 | PR221 Δ*recA*::scar | This study |

| Plasmid | Vector | Genotype | Source |
| --- | --- | --- | --- |
| pZA31R |  | Vector control | (27) |
| pZAPR1 | pZA31R | P<sub>tet</sub>-*dicF* | This study |
| pZAPR2 | pZA31R | P<sub>tet</sub>-*ypjJ* | This study |
| pZAPR3 | pZA31R | P<sub>tet</sub>-*ykfH* | This study |
| pZAPR4 | pZA31R | P<sub>tet</sub>-*yeeT* | This study |
| pZAPR5 | pZA31R | P<sub>tet</sub>-*rem* | This study |
| pZAPR6 | pZA31R | P<sub>tet</sub>-*rem* (native RBS) | This study |
| pBRPR7 | pBRCS12 | P<sub>lac</sub>-*rem* | This study |

**Table 3.**
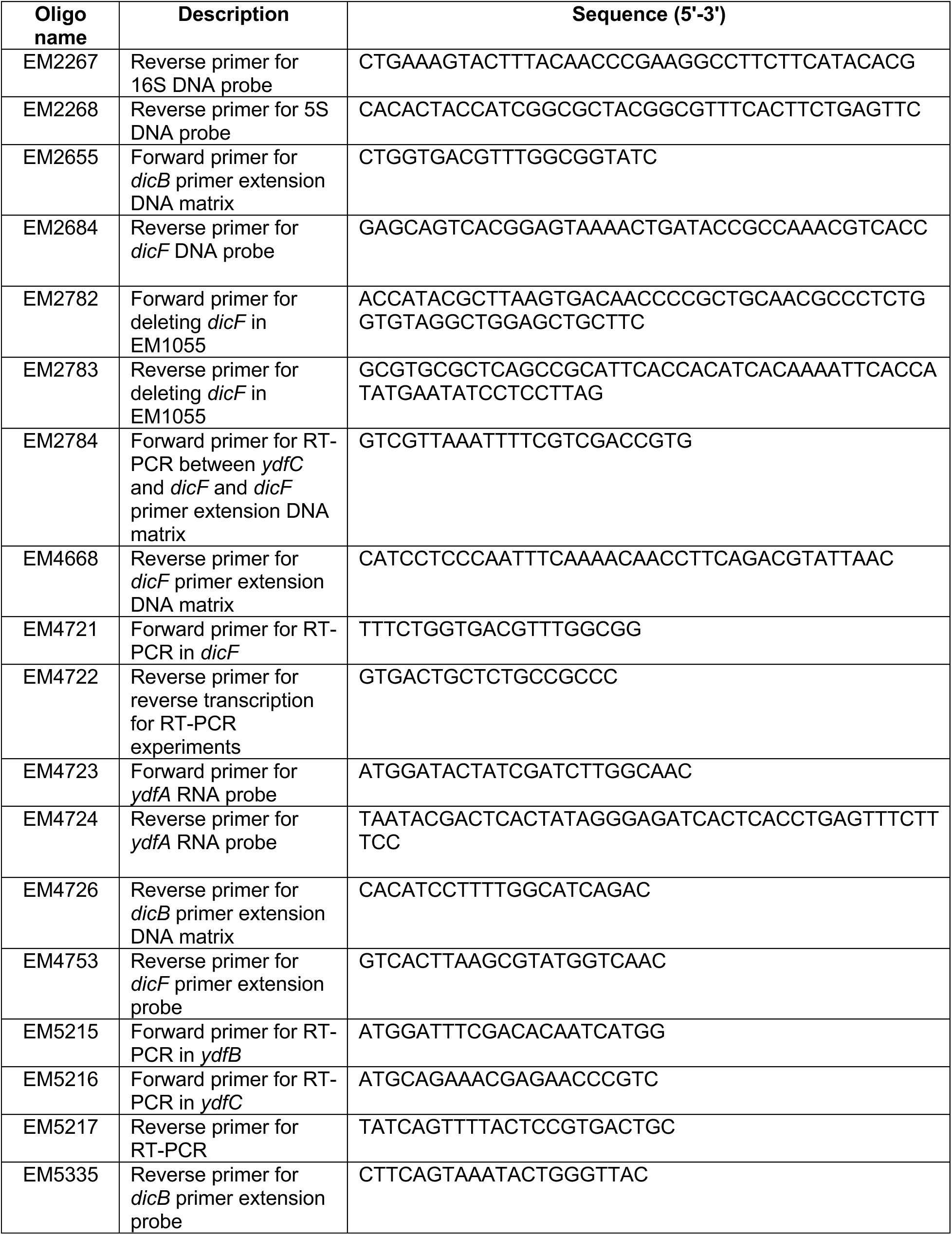
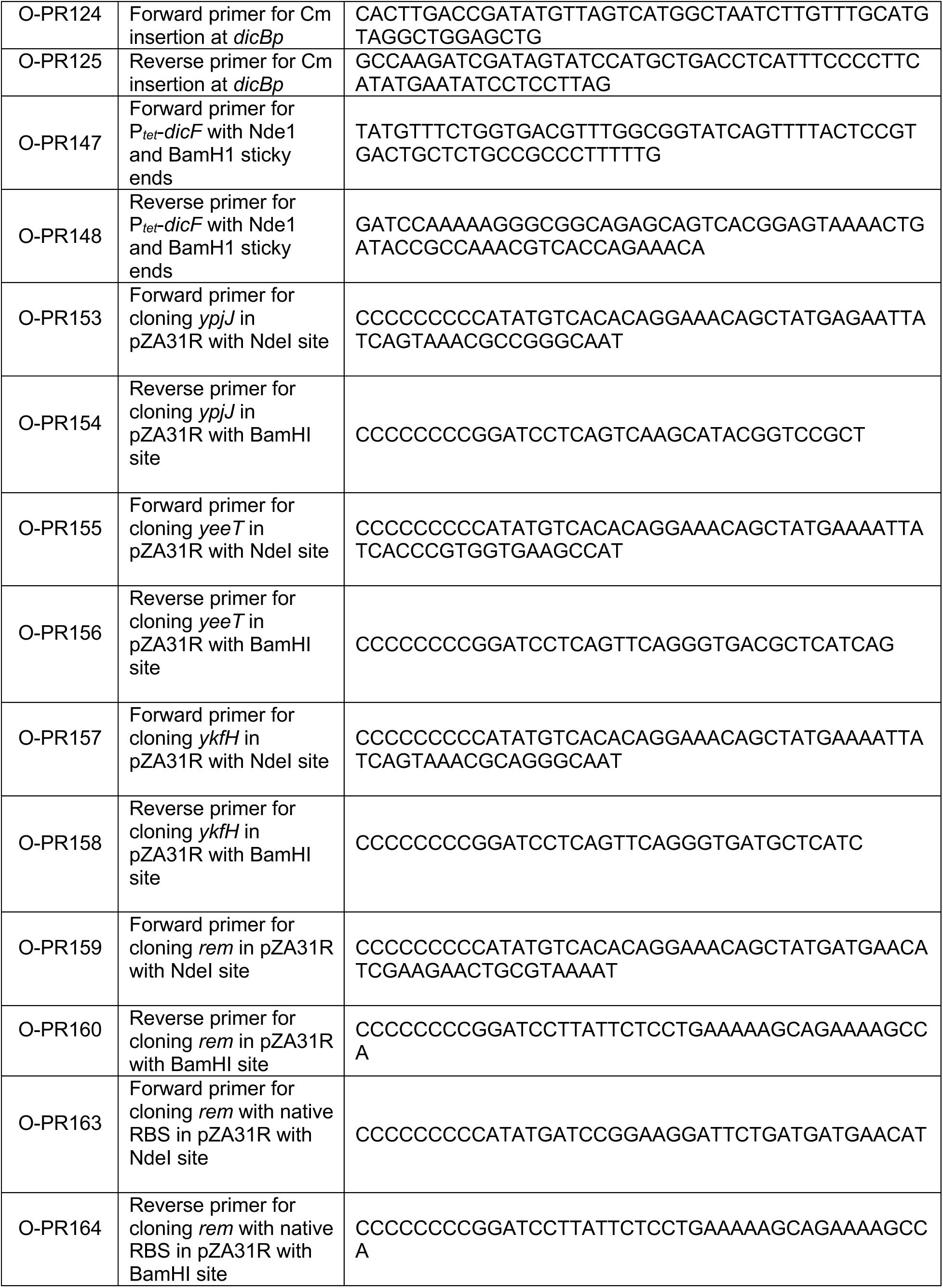
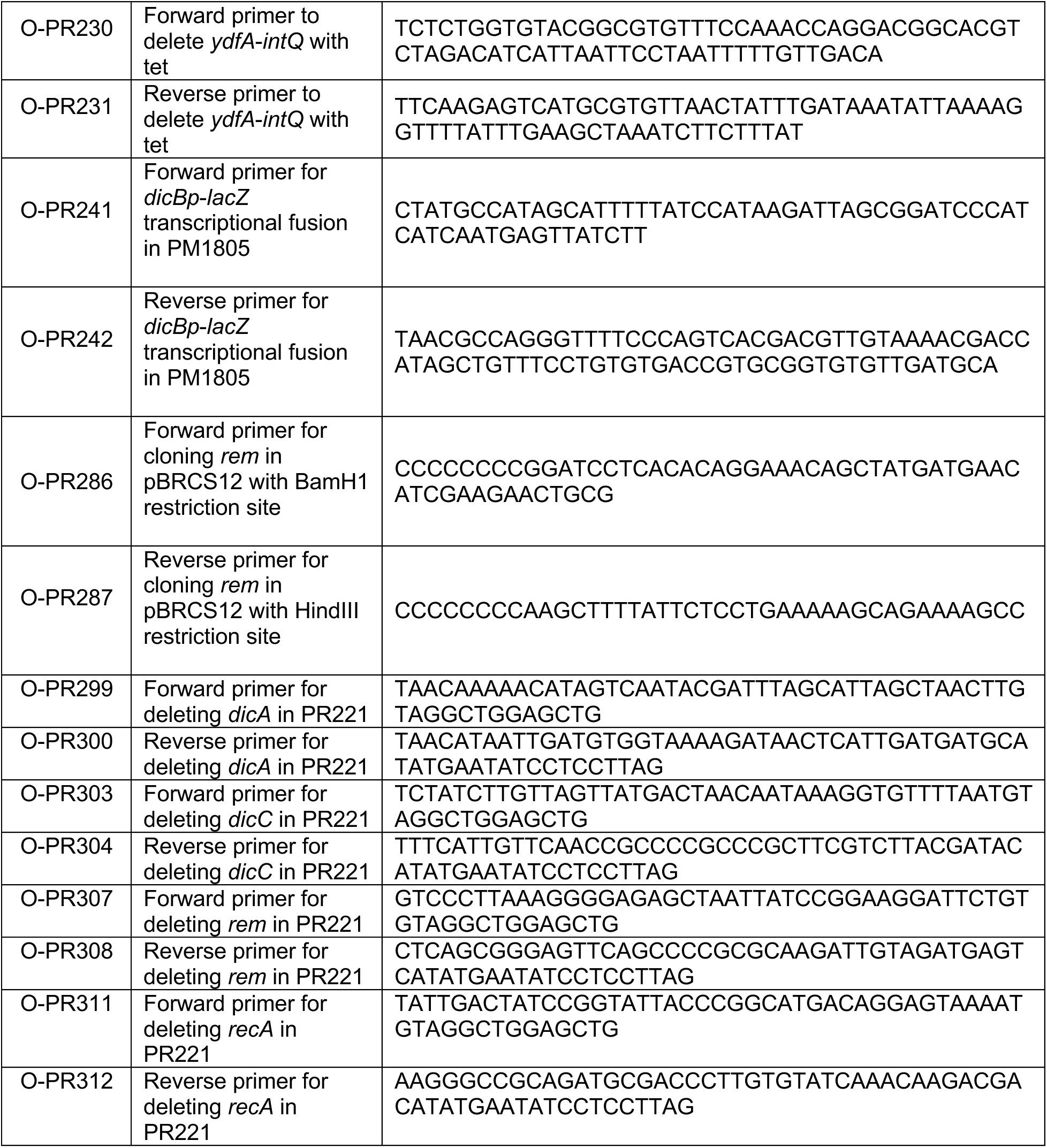
Oligonucleotides used in this study.

The in-locus *lacZ* fusion to *dicBp* was constructed as described in (18). Briefly, the primers O-PR124 and O-PR125, which contain 40 nt homology upstream and downstream of *dicBp* and 20nts homology to the FRT-chloramphenicol cassette, was amplified using pKD3 as template. This DNA fragment was recombined into the bacterial chromosome at the *dicBp* site using λ red recombination. Using PCP20, the chloramphenicol cassette was flipped out and *lacZ* was inserted using pKG136 plasmid (derivative of pCE36). Using P1*vir*, the in-locus *dicBp*-*lacZ* fusion was moved into DB166 to generate PR136.

The out-of-locus *dicBp* transcriptional fusion carried in strain PR219, was first constructed in PM1805 (29) by λ red recombination of a DNA fragment generated by amplification using primers O-PR241 and O-PR242 with WT DNA as template. This fragment contained *dicBp* and homologies to the region upstream of P_BAD_ and *lacZ*, such that P_BAD_ is replaced upon recombination. The *ydfA*-*intQ*::tet deletion was constructed by amplification using primers O-PR230 and O-PR231 and genomic DNA from strain DB181 as template to amplify the tetracycline resistance marker and recombined into the bacterial chromosome. Using P1*vir*, this deletion was moved into PR219 generating strain PR221. Strains PR226 (O-PR299/O-PR300), PR227 (O-PR303/ O-PR304), XM249 (O-PR307/ O-PR308) and PR221 Δ*recA*::cm (O-PR311/ O-PR312) were constructed using the indicated primers (names in parenthesis) with pKD3 as a template and recombined into PR221 pSIM6 by λ red recombination. The chloramphenicol cassette in PR226, PR227, XM249, and PR221 Δ*recA*::cm was removed using pCP20 leaving an FRT scar in place and generating strains PR231,PR232, XM252, and XM260, respectively.

The Δ*qin*::kan, Δ*dicB*::kan, and Δ*dicF*::kan deletions were moved from DB237, DB241, and DB252 (4) into DB166 using P1*vir* transduction to generate PR142, PR143, and PR144, respectively. Using pCP20, the kanamycin marker was removed, leaving an FRT scar resulting in strains PR145, PR146, and PR147. A Δ*dicB*::kan mutation was constructed using primers O-DB508 and O-DB509 (4) with pKD13 as the DNA template and recombined into PR147 pSIM6 to generate PR148. Using pCP20, DB166 Δ*dicF* Δ*dicB* was generated from PR148 and named PR149.

A Δ*dicF*::kan allele was constructed using primers EM2782 and EM2783 with pKD4 as the template. The resulting product was recombined in strain EM1055 (31) pKD46 to generate ENG63. Similarly, a Δ*rem*::cat deletion was constructed using primers O-PR307 and O-PR308 and recombined into EM1055 pKD46 generating ENG262.

Oligonucleotides containing NdeI and BamHI restriction sites were used to amplify the predicted antirepressor coding sequences from the chromosome of *E*. *coli* K12. The vector pZA31R (27) and PCR products were digested with NdeI and BamHI restriction enzymes and ligated using DNA ligase to generate plasmids pZAPR2, pZAPR3, pZAPR4 and pZAPR5 with the antirepressor genes under P_LtetO-1_ control. The antirepressor encoding *rem* was also cloned into pZA31R using oligonucleotides that preserved its native RBS yielding plasmid pZAPR6. DicF was cloned into pZA31R by amplification with primers O-PR147 and O-PR148 that contained the NdeI and BamH1 sites yielding pZAPR1. The vector pBRCS12 and PCR products were digested by BamH1 and HindIII restriction enzymes and ligated to generate plasmid pBRPR7 with *rem* under P*_lac_* control.

### Bioinformatic prediction of antirepressors of DicA

Using the protein sequence of GfoA, the antirepressor of Gifsy-1 prophage repressor (26), PSI-BLAST search was performed to find similar proteins in *E*. *coli*. Since many protein hits generated in the first round of PSI BLAST did not have identifiable homologs in *E*. *coli* K12, a subsequent homology search was performed using protein hits that had comparable length to GfoA. When the protein WP_171885951.1 generated from round one of PSI BLAST was used as a query, Rem encoded by Qin prophage was identified with 33% identity. Using a similar procedure with FsoA, the antirepressor of Fels-1 prophage (26), Rem was identified again as a possible candidate. By altering the BLASTp parameters, proteins YpjJ of CP4-57 prophage, YeeT of Cp4-44 prophage, and YkfH of CP4-6 prophage were also identified as other potential antirepressors in *E*. *coli* K12 when FsoA was used as a query.

### β-galactosidase assay

Strains were diluted 1:100 from overnight cultures in TB medium and grown to mid-logarithmic phase at 37°C on the rotary shaker. TB medium was supplemented with 100 µg/ml Ampicillin for pBRCS12 derivative plasmids and 0.1 mM IPTG was added to induce expression of genes under P*_lac_* control for one hour, after cells reached OD 0.2-0.3. 25 µg/ml chloramphenicol was added to TB medium when pZA31R-derived plasmids were used and 10 ng/ml anhydrous tetracycline was added to induce expression of genes under P*_tet_* control for three hours. After cultures reach the mid-logarithmic phase, β-galactosidase activity was quantified as described previously using Miller assays (34).

For experiments where β-galactosidase activity was assayed at different temperatures, strains were grown overnight at 30°C in TB medium and sub-cultured 1:100 into TB media. The strains were grown at three different temperatures, 30°C, 37°C, and 42°C. When cultures reached an OD_600_ of ∼0.8, 500 µl of sample was taken from each culture and β-galactosidase activity was quantified.

### Microscopy

Strains harboring the indicated plasmids were sub-cultured 1:100 in LB medium with 25 µg/ml chloramphenicol and 100 ng/ml anhydrous tetracycline and grown in a rotary shaker for three hours at 37°C. After three hours, the cultures were transferred to ice. 5 µl of samples were placed on a 24-by-50-mm no. 1.5 coverslip and a 1.5% agarose gel pad was placed on the cells for immobilization. Cells were then imaged using ZEISS apotome microscope under bright field setting and imaged at 40x magnification.

### Growth assays

Strains harboring the indicated plasmids were streaked on LB plates with 100 ng/ml anhydrous tetracycline to induce expression of genes under P*_tet_* control and the plates were incubated overnight at 37°C.

### Dilution plating assay

Overnight cultures of the indicated strains in Fig. 6A were serially diluted in 1x PBS. 50 µl of the 10^-5^ dilution was plated on seven LB X-Gal plates for each strain to obtain countable colonies. The plates were incubated overnight at 37°C. The next day, the blue and white colonies were counted. For each biological replicate, a total of 800 to 1600 colonies were counted from seven plates for each strain. The frequency of blue colonies was calculated using (number of blue colonies/total number of blue and white colonies)*100.

### Biolog assays

This experiment was conducted according to the protocol indicated in (10) with a key variation of adding X-Gal at a concentration of 1.25 mM to IF-0 and IF-10 growth media, instead of the dye mix. Briefly, an overnight culture of PR221 grown in LB medium was centrifuged to remove the LB medium and the pellet was resuspended in IF-0 medium. This was further diluted in IF-0 and IF-10 media supplemented with 1.25 mM X-Gal to attain the prescribed OD values stated in (10). 100 μl/well was aliquoted into plates PM 1, 2, 9 to 20 and the plates were incubated overnight at 37°C. PM1 and 2 contained different carbon sources, PM9 contained osmolytes, PM10 contained wells of different pH and PM11-20 contained different chemicals. Next day, the wells were scored based on the presence of dark blue color.

### Growth on urea and different temperatures

LB agar plates with X-Gal (40 µg/ml) or X-Gal and urea at a final concentration of 0.5%, 1%, and 2%were prepared. A single colony for each of the indicated strains was picked and streaked, and the plates were incubated overnight at 37°C. Similarly, the strains were streaked on four LB agar plates with only X-Gal (40 µg/ml) and incubated at 30, 37, 39, and 42°C overnight.

### RNA extraction and northern blot

Strains were diluted 1:1000 in LB medium or adjusted to OD 2 in M63 minimal medium supplemented with 0.2% glucose and 1µM FeSO_4_ and grown in rotary shaker. RNA samples were extracted using the classic hot-phenol protocol as described previously (1) at indicated times or OD. 5-10 µg of total RNA were loaded on a 5-10% acrylamide gel containing 8M urea. Electrotransfer was done on Hybond-XL membranes for 1h at 200 mA, then crosslinked under 254 nm UV for 45 seconds. Membranes were pre-hybridized in Church buffer for 1h at 42°C and radiolabeled DNA or RNA probes detailed in Table 3 were added overnight. Membranes were washed then exposed to phosphor screens and revealed with GE Healthcare Typhoon Trio.

### Primer extension assays

Strains were diluted 1:1000 in LB medium grown in rotary shaker. RNA samples were extracted using the classic hot-phenol chloroform protocol as described previously (1) after cells reached OD 0.5-0.6. Primer extension was then performed following the protocol previously described (8). 20 µg of total RNA was used with radiolabeled primer EM4753 to generate cDNA that was migrated on 8% acrylamide gel containing 8M urea. The sequencing ladder was generated by PCR with radiolabeled probe EM4753 from DNA matrix using primers EM2784 and EM4668. The gel was exposed on phosphor screen and revealed with GE Healthcare Typhoon Trio.

### RT-PCR assays

Cultures were adjusted to OD 2 in M63 minimal media supplemented with 0.2% glucose and 1µM FeSO_4_ and grown in a rotary shaker. RNA samples were extracted using the classic hot-phenol protocol as described previously (1) after cells reached OD 1.8. RNA extracts were treated with TURBO DNase (ThermoFischer Scientific) and reverse transcription was performed with primer EM4722, Protoscript II (NEB) and 0.1M DTT. PCR was then performed on cDNA and EM1055 strain with Taq enzyme and primers listed in Table 3.

## Acknowledgements

We thank M. Shafi Azam for his help with the sequence analyses to identify the Rem antirepressor. We thank the members of PTR’s thesis committee, James Slauch, James Imlay and Gary Olsen, for their advice throughout the course of this project. We also thank Sandy Pernitzsch from Scigraphix for graphic design of the model figure. Finally, we thank the former and current Vanderpool and Slauch laboratory members for thought-provoking discussions. This work was funded by an operating grant BMB 389354 from the Canadian Institutes of Health Research (CIHR) to EM and National Institutes of Health grants R01 GM092830 and R35 GM139557 to CKV.

---

**Figure S1.**
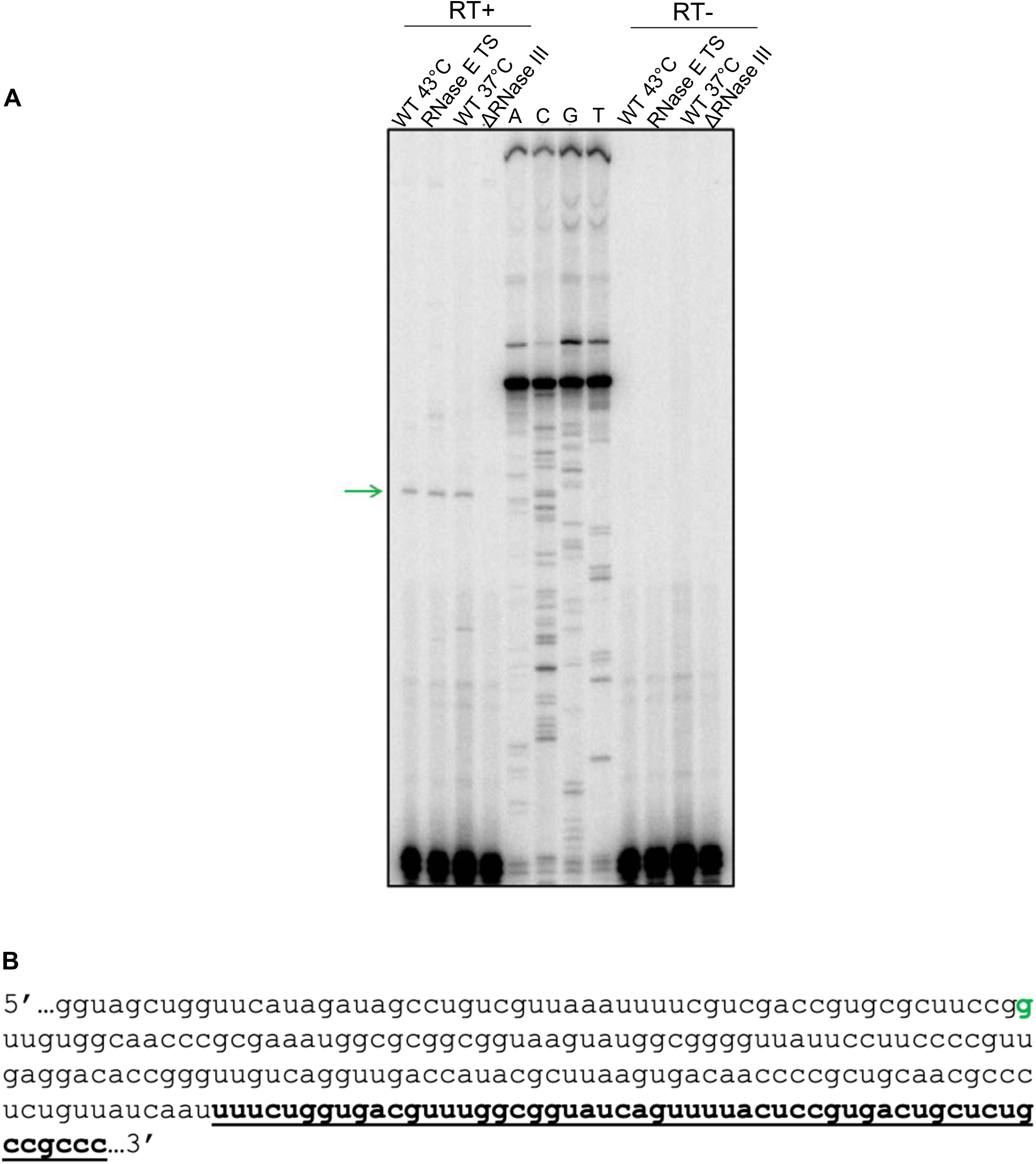
5’ end of the 190-nt DicF fragment is generated by RNase III processing of the *dicBF* transcript. A) Primer extension using radiolabeled *dicF* probe. The green arrow shows the RNase III cleavage site. B) Letter in green corresponds to the RNase III cleavage site on the DicF RNA sequence that was mapped on the primer extension in A. The 53-nt short version of DicF is underlined and in bold letters.

**Figure S2.**
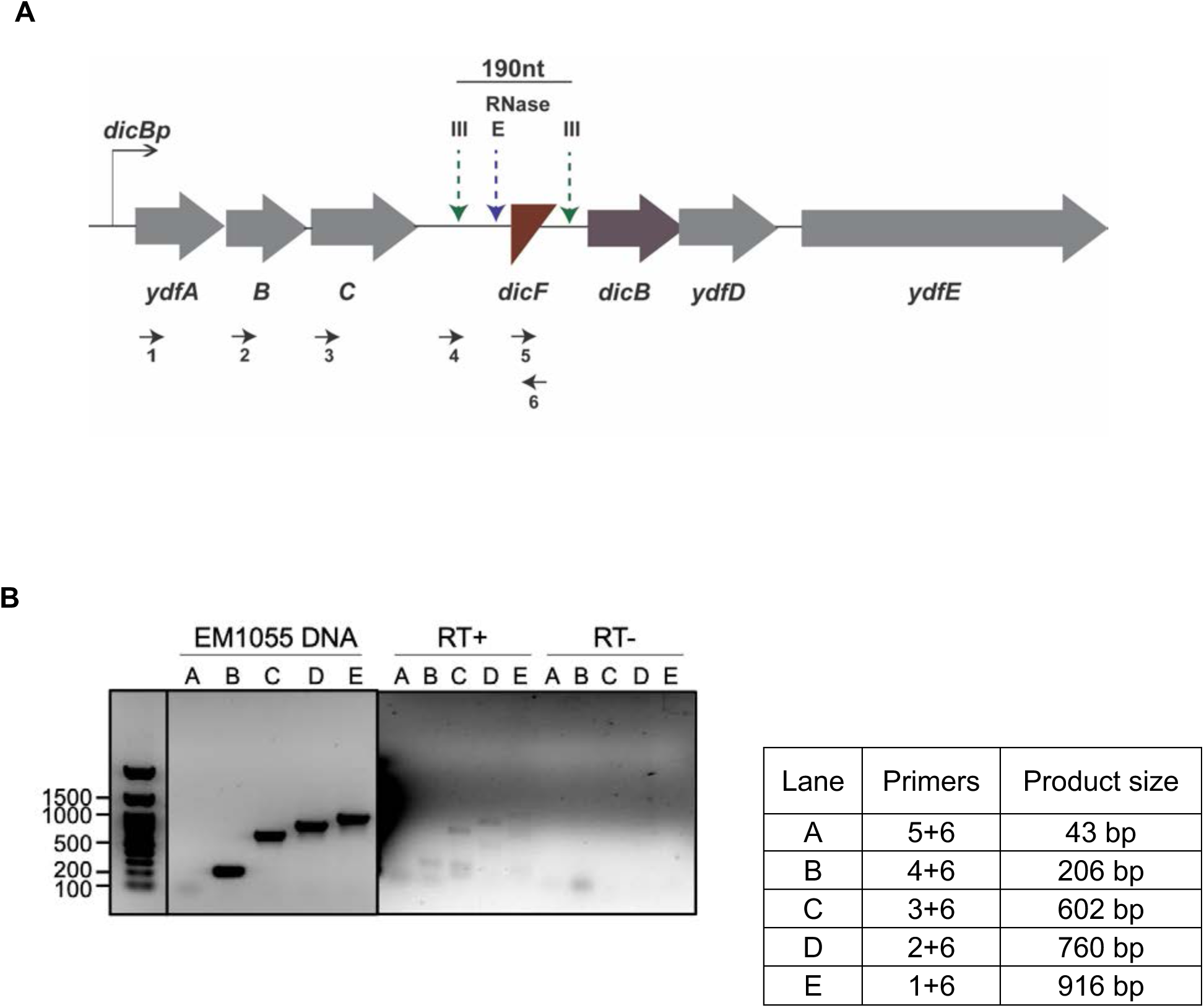
DicF is processed from the transcript starting at *dicBp*. A) Schematic representation of primers used for RT-PCR on *ydfA* to *dicF* transcript. (B) RT-PCR on *ydfA* to *dicF* transcript was carried out on RNA extracted from WT *E. coli* K12 MG1655 cells grown aerobically in M63 minimal medium supplemented with glucose and 1µM FeSO_4_. Genomic DNA of WT strain EM1055 was used as control.

**Figure S3.**
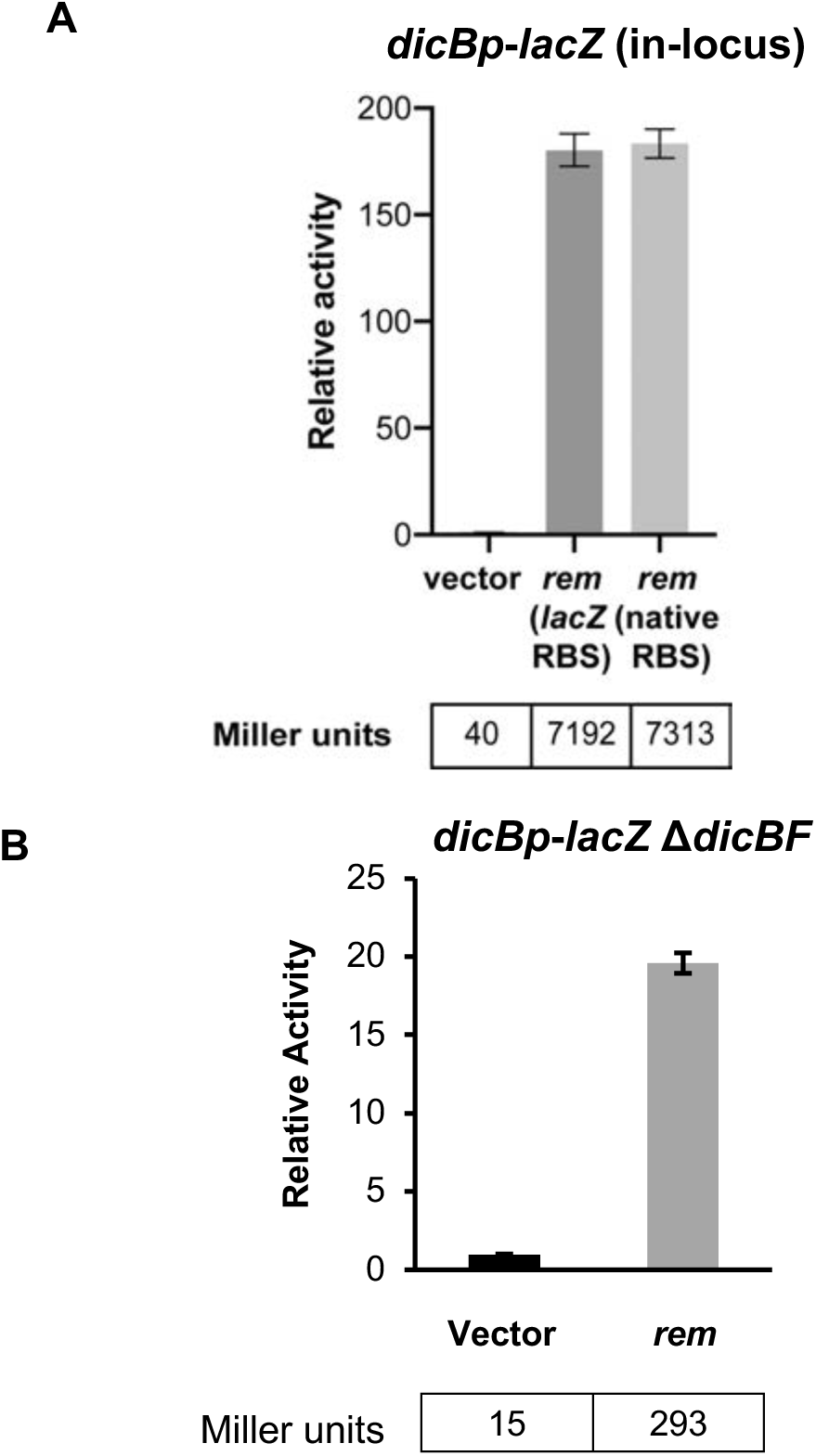
Rem is the antirepressor of *dicBp*. A) Miller assay was carried out with a *dicBp*-*lacZ* in-locus transcriptional fusion strain harboring either P*_tet_*-vector, P*_tet_*-*rem* having *lacZ* RBS, or P*_tet_*-*rem* with its native RBS. The genes were induced for three hours with 10 ng/ml anhydrous tetracycline and β-galactosidase activity was assayed B) PR221 (*dicBp*-*lacZ* Δ*dicBF*) harboring P*_lac_*-vector and P*_lac_*-*rem* was grown until early log phase, induced with 0.1 mM IPTG for one hour, and β-galactosidase activity was assayed. The relative activity was calculated by dividing the Miller units of the specific strain to that of vector control. Error bars were calculated as standard deviation from three biological replicates.

**Figure S4.**
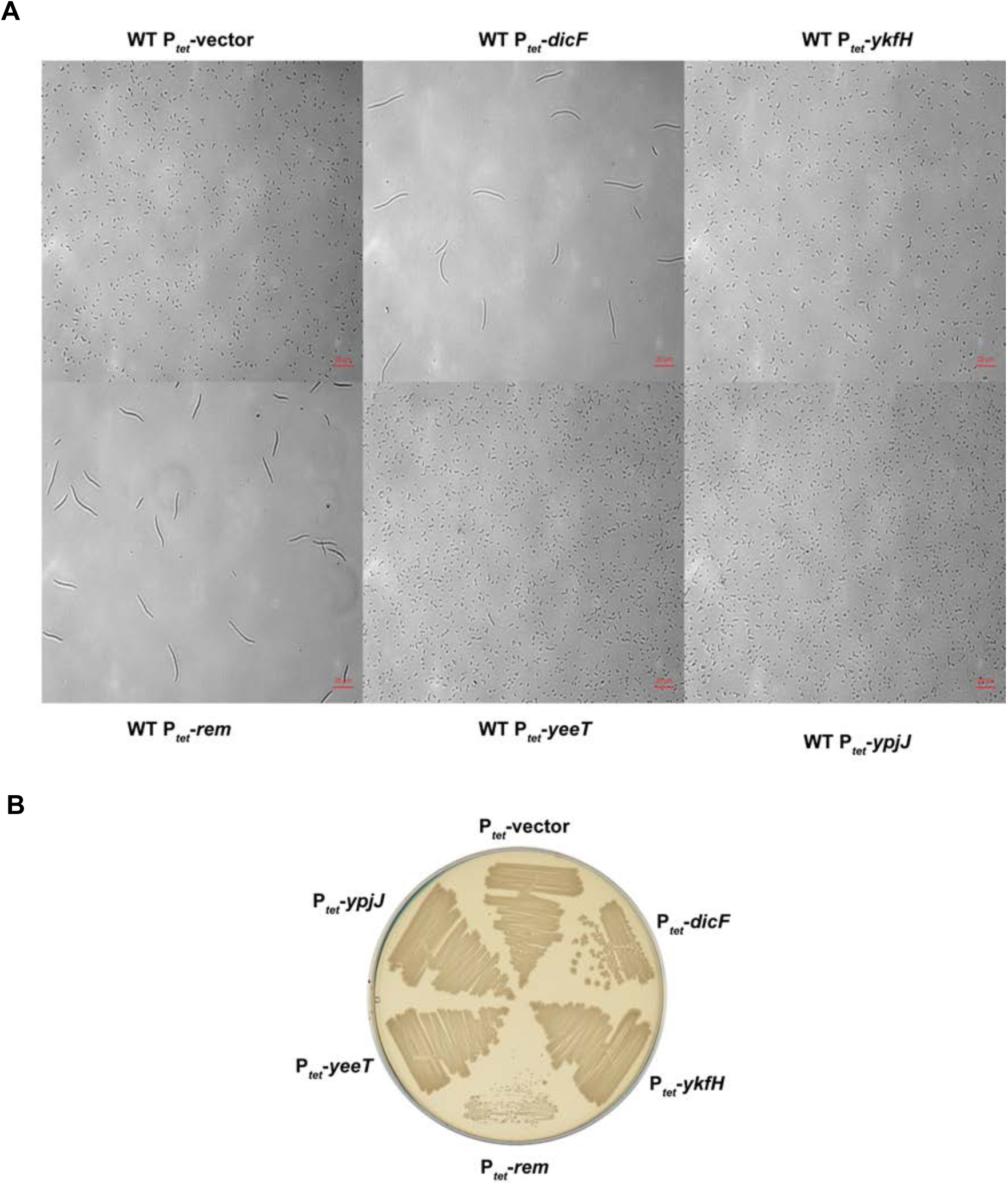
Rem is the only predicted antirepressors to cause filamentation and growth inhibition of cells. A) WT strain harboring P*_tet_*-vector, P*_tet_*-*dicF*, P*_tet_*-*ykfH*, P*_tet_*-*rem*, P*_tet_*-*yeeT* or P*_tet_*-*ypjJ* were grown for three hours in LB medium with 100 ng/ml anhydrous tetracycline and imaged using a bright field microscope. P*_tet_*-*dicF* was used as a positive control for filamentation as the sRNA DicF is known to induce filamentation (4). B) The strains from A were streaked on LB agar plates with 100 ng/ml anhydrous tetracycline and incubated overnight at 37°C. P*_tet_*-*dicF* was used as a positive control for growth inhibition as prolonged expression of DicF is toxic to the cells (4).

**Figure S5.**
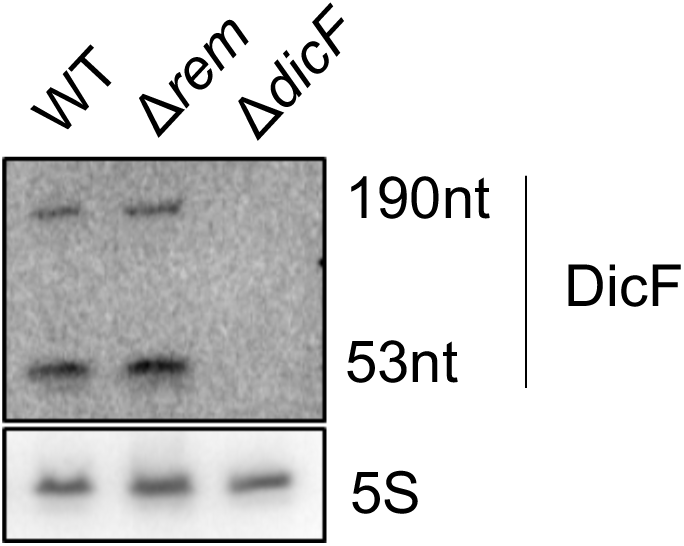
DicF expression during growth in M63 minimal media is independent of Rem. Northern blot showing the expression of DicF in a *rem* mutant strain. RNA was extracted at OD_600_ of 1.7-1.8 in M63 minimal media supplemented with 0.2% glucose and 1µM FeSO4. 5S RNA was used as a loading control.

## References

1. Aiba H, Adhya S, de Crombrugghe B. 1981. Evidence for two functional gal promoters in intact Escherichia coli cells. J. Biol. Chem. 256(22):11905–10

2. Amemiya HM, Goss TJ, Nye TM, Hurto RL, Simmons LA, Freddolino PL. 2022. Distinct heterochromatin-like domains promote transcriptional memory and silence parasitic genetic elements in bacteria. EMBO J. 41(3):e108708

3. Azam MS, Vanderpool CK. 2017. Translational regulation by bacterial small RNAs via an unusual Hfq-dependent mechanism. Nucleic Acids Res. 46(5):2585–99

4. Balasubramanian D, Ragunathan PT, Fei J, Vanderpool CK. 2016. A prophage-encoded small RNA controls metabolism and cell division in *Escherichia coli*. mSystems. 1(1):e00021–15

5. Béjar S, Bouché JP. 1985. A new dispensable genetic locus of the terminus region involved in control of cell division in *Escherichia coli*. Mol. Gen. Genet. 201(2):146–50

6. Béjar S, Bouché F, Bouché JP. 1988. Cell division inhibition gene *dicB* is regulated by a locus similar to lamboid bacteriophage immunity loci. Mol. Gen. Genet. 212(1):11–19

7. Béjar S, Cam K, Bouché JP. 1986. Control of cell division in *Escherichia coli*. DNA sequence of dicA and of a second gene complementing mutation dicA1, dicC. Nucleic Acids Res. 14(17):6821–33

8. Beroual W, Prévost K, Lalaouna D, Ben Zaina N, Valette O, et al. 2022. The noncoding RNA CcnA modulates the master cell cycle regulators CtrA and GcrA in Caulobacter crescentus. PLOS Biol. 20(2):e3001528

9. Bobay LM, Touchon M, Rocha, EPC. 2014. Pervasive domestication of defective prophages by bacteria. Proc Natl Acad Sci U S A. 111(33):12127–32

10. Bochner BR. 2001. Phenotype MicroArrays for High-Throughput Phenotypic Testing and Assay of Gene Function. Genome Res. 11(7):1246–55

11. Bouché F, Bouché JP. 1989. Genetic evidence that DicF, a second division inhibitor encoded by the *Escherichia coli dicB* operon, is probably RNA. Mol. Microbiol. 3(7):991– 94

12. Cam K, Béjar S, Gil D, Bouché JP. 1988. Identification and sequence of gene *dicB*: translation of the division inhibitor from an in-phase internal start. Nucleic Acids Res. 16(14A):6327–38

13. Campbell AM. 1998. Prophages and Cryptic Prophages. In Bacterial Genomes, ed FJ de Bruijn, JR Lupski, GM Weinstock, pp. 23–29. Boston, MA: Springer

14. Casjens S. 2003. Prophages and bacterial genomics: what have we learned so far? Mol. Microbiol. 49(2):277–300

15. Casjens SR, Hendrix RW. 2015. Bacteriophage lambda: Early pioneer and still relevant. Virology. 479–480:310–30

16. Chan W, Costantino N, Li R, Lee SC, Su Q, Melvin D, Court DL, Liu P. 2007. A recombineering based approach for high-throughput conditional knockout targeting vector construction. Nucleic Acids Res. 35(8):e64

17. Datsenko KA, Wanner BL. 2000. One-step inactivation of chromosomal genes in *Escherichia coli* K-12 using PCR products. Proc Natl Acad Sci U S A. 97(12):6640–45

18. Ellermeier CD, Janakiraman A, Slauch JM. 2002. Construction of targeted single copy *lac* fusions using λ Red and FLP-mediated site-specific recombination in bacteria. Gene. 290(1–2):153–61

19. Faubladier M, Cam K, Bouché JP. 1990. *Escherichia coli* cell division inhibitor DicF-RNA of the dicB operon. Evidence for its generation in vivo by transcription termination and by RNase III and RNase E-dependent processing. J. Mol. Biol. 212(3):461–71

20. Fortier LC, Sekulovic O. 2013. Importance of prophages to evolution and virulence of bacterial pathogens. Virulence. 4(5):354–65

21. Gerdes K, Bech FW, Jørgensen ST, Løbner-Olesen A, Rasmussen PB, et al. 1986. Mechanism of postsegregational killing by the hok gene product of the parB system of plasmid R1 and its homology with the relF gene product of the E. coli relB operon. EMBO J. 5(8):2023–29

22. Gimble FS, Sauer RT. 1985. Mutations in bacteriophage lambda repressor that prevent RecA-mediated cleavage. J. Bacteriol. 162(1):147–54

23. Harrison E, Brockhurst MA. 2017. Ecological and evolutionary benefits of temperate phage: what does or doesn’t kill you makes you stronger. BioEssays. 39(12):1700112

24. Johnson JE, Lackner LL, de Boer PAJ. 2002. Targeting of (D)MinC/MinD and (D)MinC/DicB complexes to septal rings in *Escherichia coli* suggests a multistep mechanism for MinC-mediated destruction of nascent FtsZ rings. J. Bacteriol. 184(11):2951–62

25. Johnson JE, Lackner LL, Hale CA, De Boer PAJ. 2004. ZipA is required for targeting of DMinC/DicB, but not DMinC/MinD, complexes to septal ring assemblies in *Escherichia coli*. J. Bacteriol. 186(8):2418–29

26. Lemire S, Figueroa-Bossi N, Bossi L. 2011. Bacteriophage Crosstalk: Coordination of Prophage Induction by Trans-Acting Antirepressors. PLoS Genet. 7(6):e1002149

27. Levine E, Zhang Z, Kuhlman T, Hwa T. 2007. Quantitative characteristics of gene regulation by small RNA. PLoS Biol. 5(9):1998–2010

28. Little JW. 1999. Robustness of a gene regulatory circuit. EMBO J. 18(15):4299–4307

29. Mandin P, Gottesman S. 2009. A genetic approach for finding small RNAs regulators of genes of interest identifies RybC as regulating the DpiA/DpiB two-component system. Mol. Microbiol. 72(3):551–65

30. Massé E, Escorcia FE, Gottesman S. 2003. Coupled degradation of a small regulatory RNA and its mRNA targets in Escherichia coli. Genes Dev. 17(19):2374–83

31. Massé E, Gottesman S. 2002. A small RNA regulates the expression of genes involved in iron metabolism in Escherichia coli. Proc. Natl. Acad. Sci. 99(7):4620–25

32. Mathieu A, Dion M, Deng L, Tremblay D, Moncaut E, Shah SA, Stokholm J, Krogfelt KA, Schjørring S, Bisgaard H, Nielsen DS, Moineau S, Petit MA. 2020. Virulent coliphages in 1-year-old children fecal samples are fewer, but more infectious than temperate coliphages. Nat. Commun. 11(1):378

33. Melson EM, Kendall MM. 2019. The sRNA DicF integrates oxygen sensing to enhance enterohemorrhagic *Escherichia coli* virulence via distinctive RNA control mechanisms. Proc Natl Acad Sci U S A. 116(28):14210–15

34. Miller J. 1972. Experiments in molecular genetics. Cold Spring Harbor Laboratory, Cold Spring Harbor, NY

35. Mount DW. 1976. A method for the isolation of phage mutants altered in their response to lysogenic induction. Mol. Gen. Genet. MGG. 145(2):165–67

36. Murashko ON, Lin-Chao S. 2017. *Escherichia coli* responds to environmental changes using enolasic degradosomes and stabilized DicF sRNA to alter cellular morphology. Proc Natl Acad Sci U S A. 114(38):E8025–34

37. Murray NE, Gann A. 2007. What has phage lambda ever done for us? Curr. Biol. 17(9):R305–12

38. Oppenheim AB, Kobiler O, Stavans J, Court DL, Adhya S. 2005. Switches in Bacteriophage Lambda Development. Annu. Rev. Genet. 39(1):409–29

39. Pan H, Dong K, Rao L, Zhao L, Wang Y, Liao X. 2019. The Association of Cell Division Regulated by DicC With the Formation of Viable but Non-culturable *Escherichia coli* O157:H7. Front. Microbiol. 10:2850

40. Ragunathan PT, Vanderpool CK. 2019. Cryptic-Prophage-Encoded Small Protein DicB Protects *Escherichia coli* from Phage Infection by Inhibiting Inner Membrane Receptor Proteins. J. Bacteriol. 201(23):JB.00475-19

41. Schubert RA, Dodd IB, Egan JB, Shearwin KE. 2007. Cro’s role in the CI Cro bistable switch is critical for ’s transition from lysogeny to lytic development. Genes & Dev. 21(19):2461–72

42. Tétart F, Bouché JP. 1992. Regulation of the expression of the cell-cycle gene *ftsZ* by DicF antisense RNA. Division does not require a fixed number of FtsZ molecules. Mol. Microbiol. 6(5):615–20

43. Toman Z, Dambly-Chaudière C, Tenenbaum L, Radman M. 1985. A system for detection of genetic and epigenetic alterations in *Escherichia coli* induced by DNA-damaging agents. J. Mol. Biol. 186(1):97–105

44. Touchon M, Bernheim A, Rocha EPC. 2016. Genetic and life-history traits associated with the distribution of prophages in bacteria. ISME J. 10(11):2744–54

45. Wang X, Kim Y, Ma Q, Hong SH, Pokusaeva K, Sturino JM, Wood TK. 2010. Cryptic prophages help bacteria cope with adverse environments. Nat. Commun. 1(1):147

46. Yu D, Ellis HM, Lee E-C, Jenkins NA, Copeland NG, Court DL. 2000. An efficient recombination system for chromosome engineering in *Escherichia coli*. Proc Natl Acad Sci U S A. 97(11):5978–83

47. Yun SH, Ji SC, Jeon HJ, Wang X, Kim SW, Bak G, Lee Y, Lim HM. 2012. The CnuK9E H-NS Complex Antagonizes DNA Binding of DicA and Leads to Temperature-Dependent Filamentous Growth in *E*. *coli*. PLoS One. 7(9):e45236

